# Characterizing the heterogeneity of neurodegenerative diseases through EEG normative modeling

**DOI:** 10.1101/2024.07.22.604542

**Authors:** Judie Tabbal, Aida Ebadi, Ahmad Mheich, Aya Kabbara, Bahar Güntekin, Görsev Yener, Veronique Paban, Ute Gschwandtner, Peter Fuhr, Marc Verin, Claudio Babiloni, Sahar Allouch, Mahmoud Hassan

## Abstract

Neurodegenerative diseases such as Parkinson’s (PD) and Alzheimer’s (AD) exhibit considerable heterogeneity of functional brain features within patient populations, complicating diagnosis, treatment, prognosis, and drug discovery. Here, we use electroencephalography (EEG) and normative modeling to investigate neurophysiological oscillatory mechanisms underpinning this heterogeneity. To this aim, we use resting-state EEG activity collected by 14 clinical units, in healthy older persons (n=499) and patients with PD (n=237) and AD (n=197), aged over 40 years old. Spectral and source connectivity analyses of EEG activity provided EEG features for normative modeling and deviation measures in the PD and AD patients. Normative models confirmed significant deviations of the EEG features in PD and AD patients over population norms, characterized by high heterogeneity and frequency-dependence. The percentage of patients with at least one deviating EEG feature was ∼30% for spectral measures and ∼80% for functional source connectivity. Notably, the spatial overlap of the deviant EEG features did not exceed 60% for spectral analysis and 25% for functional source connectivity analysis. Furthermore, the patient-specific deviations were correlated with relevant clinical measures, such as the UPDRS for PD (*⍴*=0.24, *p*=0.025) and the MMSE for AD (*⍴*=-0.26, *p*=0.01), indicating that greater deviations from normative EEG features are associated with worsening score values. These results suggest that the deviation percentage from EEG norms may enrich clinical assessment in PD and AD patients at individual levels in the framework of Precision Neurology.

## Introduction

As the global population ages rapidly, the number of people over sixty is expected to double by 2050^1^. This trend coincides with an expected rise in the prevalence of patients with age-related progressive neurodegenerative diseases, such as Parkinson’s (PD) and Alzheimer’s (AD) diseases, belonging to cognitive deficits and disabilities in the activities of daily living, culminating in the diagnosis of dementia. Current estimates project that the number of people living with dementia is about 50 million and will triple to 153 million by 2050^2^. Among them, the majority suffer from AD. Furthermore, the number of people with PD is expected to reach around 16 million globally^3^. Notably, there has been no therapy able to cure or stop AD and PD until now, motivating more research on age-related brain changes in pathological aging.

Over the past three decades, advances in neuroimaging and electroencephalographic (EEG) techniques have provided invaluable biomarkers reflecting brain dysfunctions, enhancing efforts to describe aging processes, distinguish between healthy and diseased individuals, and predict disease progression (Vecchio et al. 2013; Dennis and Thompson 2014; Li et al. 2015; MacDonald and Pike 2021; Pini et al. 2016; Ryman and Poston 2020; Frisoni et al. 2010; Talwar et al. 2021). However, most research studies have traditionally focused on MRI and EEG biomarkers derived from group analyses, comparing healthy controls to patient populations or patient populations along the disease course or in response to treatments (Verdi et al., 2021). However, by averaging data across a population, group analyses tend to obscure individual variations, which can mask important differences in disease progression and response to treatment at individual level ls (Marquand et al. 2016; Verdi et al. 2021). According to Precision Medicine, this ‘*one-size-fits-all’* methodology fails to account for the unique genetic, environmental, and lifestyle factors that influence each patient’s condition. Consequently, treatments based on group data may be less effective or even inappropriate for certain individuals, leading to suboptimal outcomes.

In this context, normative models emerge as a powerful tool to make individual-level inferences by describing the population norm and then assessing the degree to which each individual deviates from these norms (Verdi et al. 2021; Marquand et al. 2016; 2019). While normative charts are well-established in other domains, such as pediatric growth charts, their application to neuroimaging data is relatively recent. Similar to how growth charts track a child’s development by comparing their height and weight to peers, recent MRI studies combined with normative models have endeavored to chart analogous trajectories for brain phenotypes (e.g., grey and white matter volumes, mean cortical thickness, total surface area, etc.) to map lifespan age-related changes in brain structure (Bethlehem et al. 2022; Rutherford et al. 2022; 2023) and characterize structural/functional heterogeneity in psychiatric disorders (Segal et al. 2023; Wolfers et al. 2020; Zabihi et al. 2020), schizophrenia and bipolar disorder (Wolfers et al. 2018), as well as in neurodegenerative disease as Alzheihmer’s disease (AD) (Verdi et al. 2023; Rutherford et al. 2022; Sun et al. 2023; Huo et al. 2024).

While recent studies have focused on MRI data, the EEG-based normative modeling field remains largely unexplored. Notably, (Lefebvre et al. 2018) used normative models to investigate variability among autistic patients as compared to healthy controls. More recently, we charted the trajectory of brain development in a population aged between 5 and 18 years and mapped the heterogeneity of psychiatric diseases as reflected in spectral power density spectra computed at scalp electrodes from resting-state eyes-closed EEG activity and cortical source functional connectivity estimated from that activity (Ebadi et al. 2024). In this study, we utilized normative models and EEG data from 14 datasets to achieve two primary objectives. First, we aimed to chart the trajectory of standard EEG features, including EEG relative power and source-level functional connectivity, in a cohort of healthy control (HC) individuals over 40 years old. Second, we leveraged individual-level inferences from the normative models to map the heterogeneity within AD and PD groups.

## Results

### Data description

Resting-state eyes-closed EEG data was collected from subjects aged 40 to 92 years, across 14 sites (<u>Materials and Methods</u>). Normative models were trained on 400 (46% M) HC participants while 99 (42% M) HCts participants were held out as a comparison group against the clinical cohort. The clinical groups comprised 237 PD patients (65% M), and 197 AD patients (37% M). An overview of the age distribution across groups, sex, and sites is provided in Fig. 1.

**Fig. 1.**
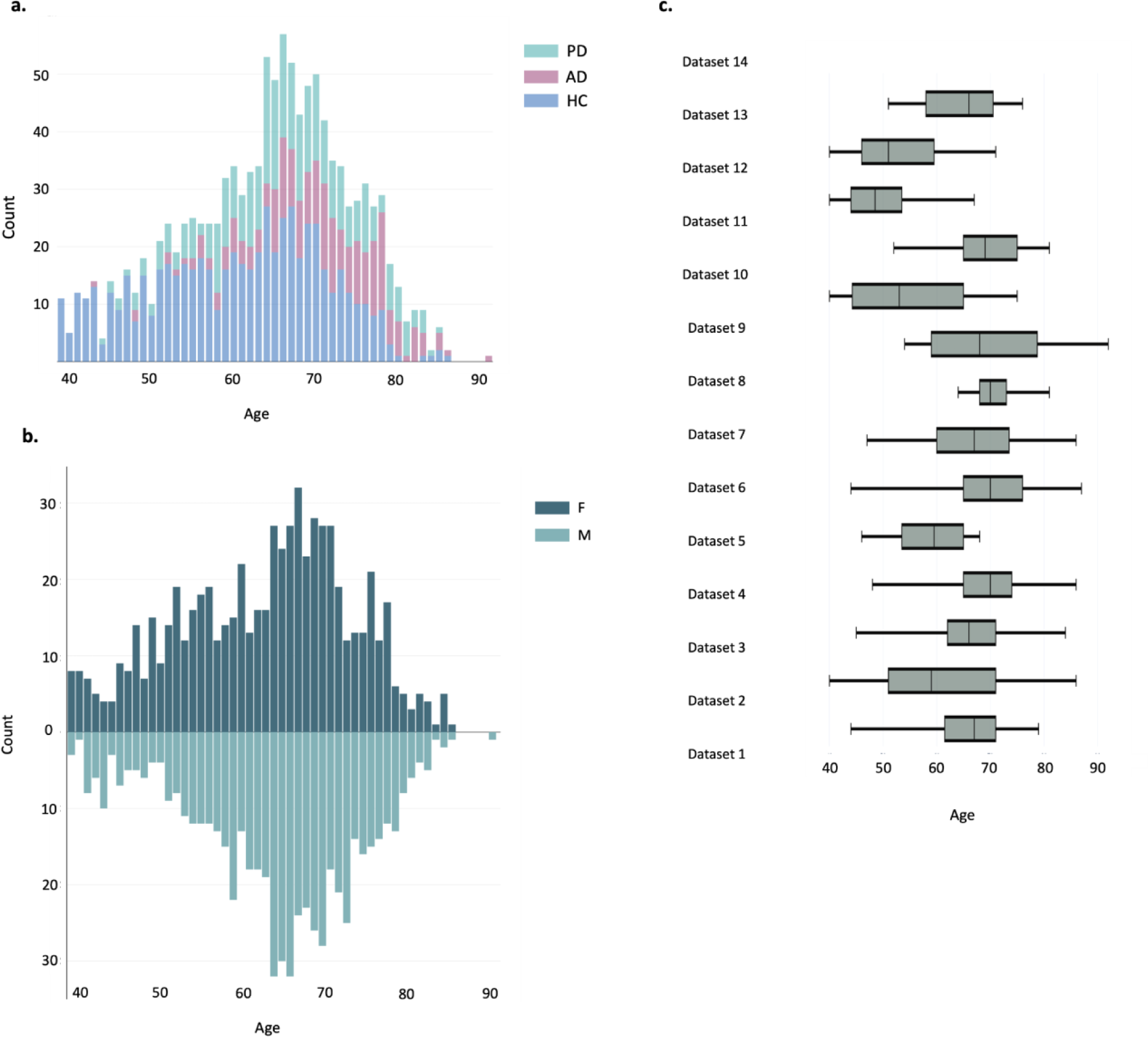
Age distribution overview for the whole study. **(a)** Age distribution across healthy and clinical groups. **(b)** Age distribution bar plots across sex for each group (lighter color for female). **(c)** Age distribution box plots across different sites.

### Building the normative model

Using a reference sample of HC individuals, we trained a GAMLSS for each EEG feature (see Methods). Specifically, relative EEG power and cortical source functional connectivity values were computed at each scalp electrode/functional connection between cortical source pairs for the frequency bands of interest (i.e., delta, theta, alpha, beta, and gamma) were modeled. The optimal distribution family, model parameters (*μ*, *σ*, *v*, *τ*), and covariates are indicated in Tables S4 and S5. The performance and robustness of our models are depicted in Fig. S3-S6. Leveraging our reference healthy cohort, we generated the typical trajectory observed in spectral and functional connectivity features. An example of these normative trajectories at the alpha band is presented in Fig. 2, showcasing the averaged spectral and connectivity features across channels and connections Results across frequency bands can be found in Fig. S7 and S8.

**Fig. 2.**
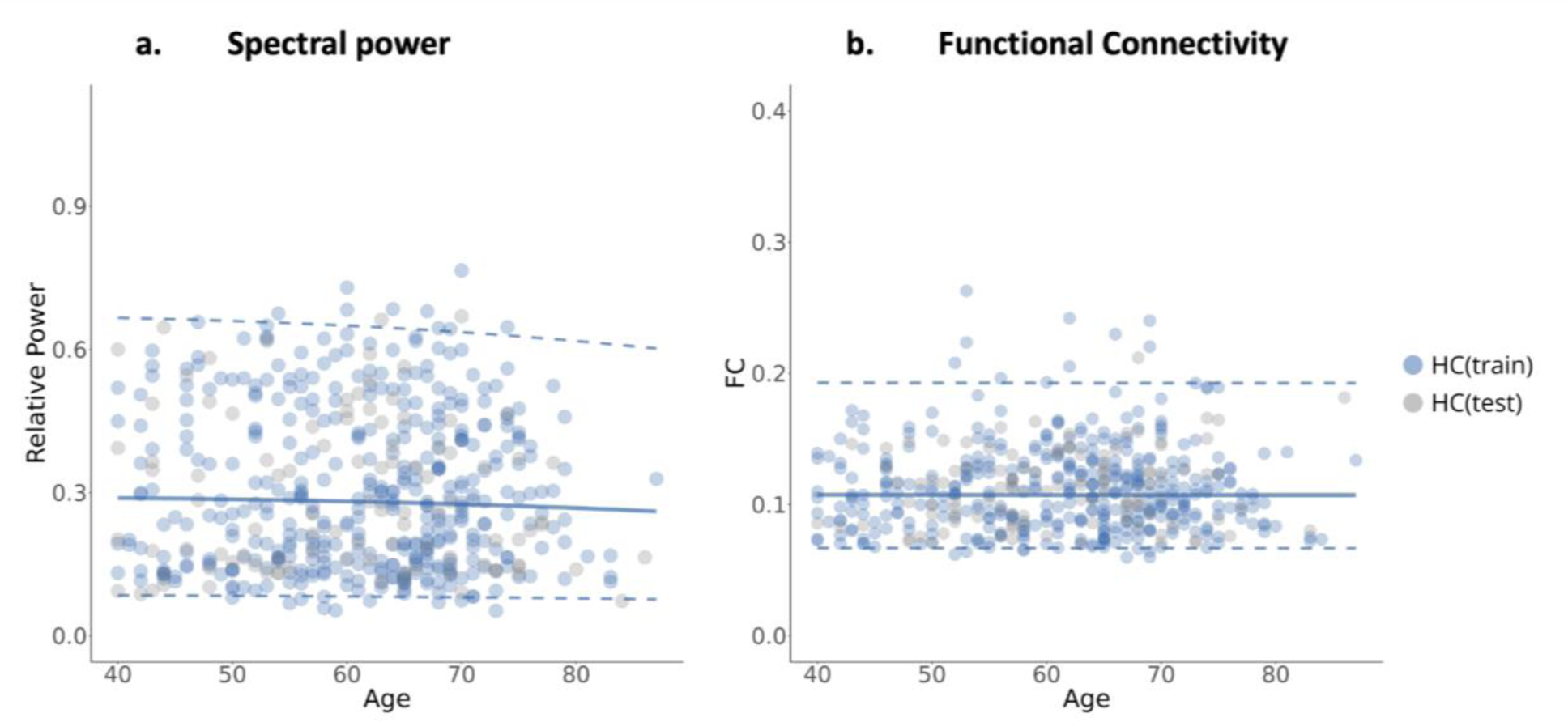
Normative aging trajectories of spectral power and functional connectivity in the alpha band. The median (50th percentile) is depicted with a solid blue line, while the 5th and 95th percentiles are indicated by dotted blue lines. **(a)** NM of the relative power averaged over all channels. **(b)** NM of FC values averaged over all connections.

### Heterogeneity within neurodegenerative diseases

After generating the normative trajectories, we projected individuals diagnosed with PD and AD, alongside an HC test group, onto these models, and derived subject-specific deviation scores. Following previous work done by (Rutherford et al. 2023; 2022; Segal et al. 2023; Verdi et al. 2023; Wolfers et al. 2018; Zabihi et al. 2020) to quantify within-group heterogeneity, we computed the number of extremely deviated channels (scalp electrodes)/functional source connections (|z-scores|>2) per participant (Fig. S9-S10) and the percentage of participants with at least one extremely deviated channel/connection (Table S6-S7).

Results of the spectral analysis showed that the percentages of participants having at least one deviation were relatively low with the highest values occurring at the theta band (PD: 31.36%, AD: 27.41%, *positive deviation*). The values of the negative deviation reached their maximum at the beta band (PD: 12.71%, AD: 23.35%, *negative deviation*), indicating that a good proportion of participants exhibited significant similarities with the HC persons forming the training HC(train) group. The number of channels (EEG power density computed at scalp electrodes) per participant negatively deviating from the normative model was significantly greater for AD compared to HC(test) at the alpha and beta bands (*p*<0.01, Mann-Whitney test). Compared to the HC individuals, these patients had lower EEG power density at those bands. Regarding positive deviations, both PD and AD groups showed significant differences compared to the HC(test) group at the theta, alpha, and beta bands (*p<0.05*). Compared to the HC group, these patients had higher EEG power density at those bands.

Unlike the findings from the EEG spectral analysis, we observed an increase in the number of participants exhibiting at least one extreme negative/positive deviation in the EEG cortical source connectivity analysis. Detailed percentages across all frequency bands and disorders are provided in Supplementary Table S7. Fig. 3 demonstrates an example of these measures at the theta band. Particularly noteworthy are the percentages of negative extreme deviations in PD across all frequency bands, where they reached 86.86% at the delta band. Interestingly, PD exhibited a greater number of extreme negative deviations compared to HC(test), while AD displayed fewer. Conversely, the PD group had significantly fewer deviated positive EEG connections at source pairs than the HC(test) group across all EEG frequency bands except delta, whereas AD exhibited a significantly higher number of deviated EEG source connections only at the alpha band (*p<0.05*).

**Fig. 3.**
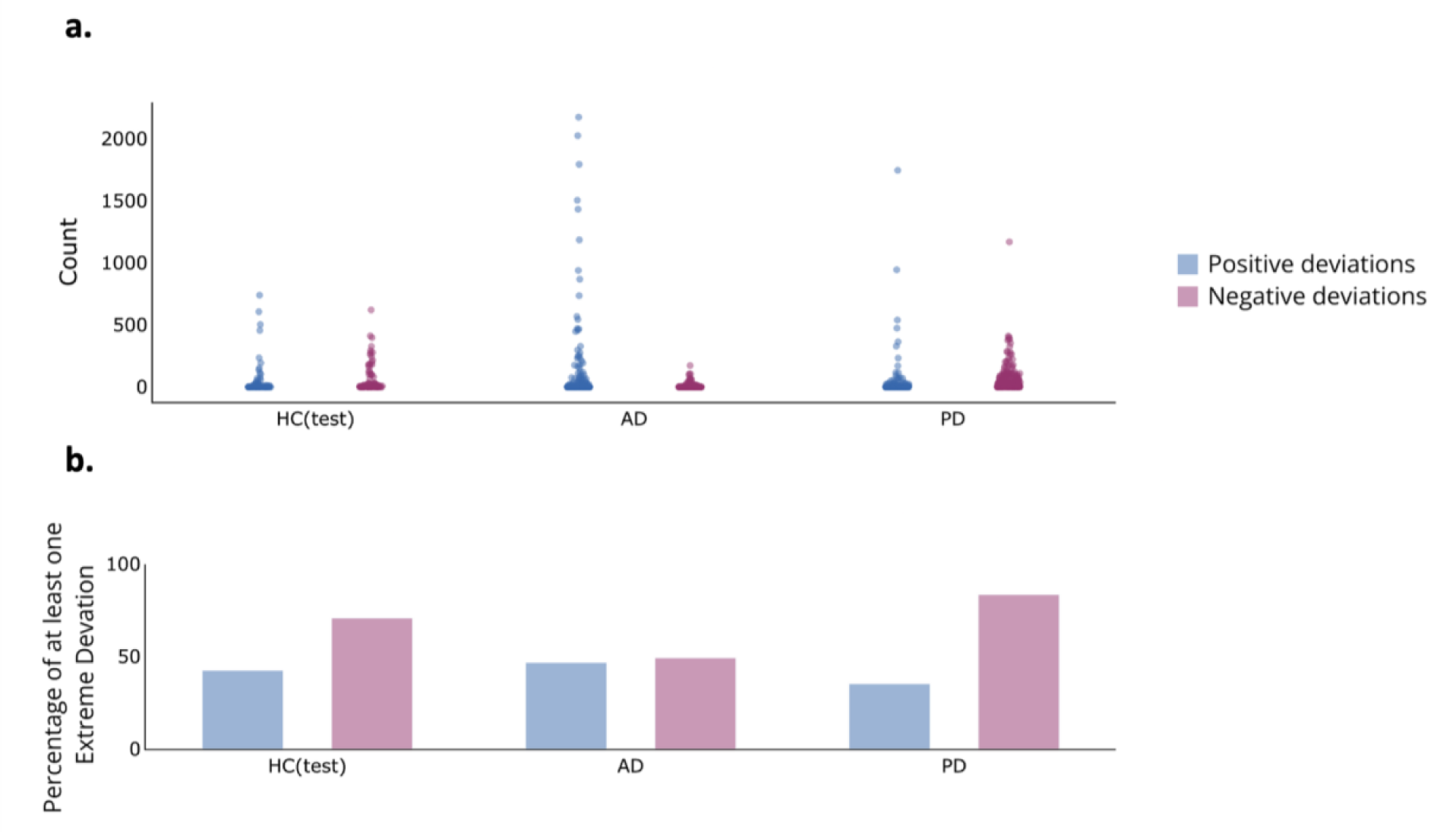
Distributions of the numbers of extremely deviated connections per subject across groups for theta band (a) and Percentages of subjects with at least one extreme deviation per group (b). Blue (Red) violins represent positive (negative) deviations.

Next, interested in searching for a common deviation pattern within each group, we evaluated the spatial overlap of extreme deviations by calculating the percentage of participants with extreme deviations (|z-score|>2) at each channel (EEG power density at scalp electrode)/ functional connection at source pairs within each group to create deviation overlap maps (Fig.4 a, b). For the EEG spectral features at scalp electrodes, a certain consistency in spatial locations among individuals within the same group was observed, more prominently within the HC(test) group than within the PD and AD groups, across all frequency bands. Specifically, we noted some deviated channels that are shared by more than 60% of individuals within the same group. These cases correspond to the HC(test) participants evaluated at the alpha and beta bands (negative deviation), as well as the delta and theta bands (positive deviation). More than 60% of the PD patients shared common extremely deviated channels at the theta band (positive deviation). Similar observations were noted in the case of AD patients at the beta band (negative deviation).

**Fig. 4.**
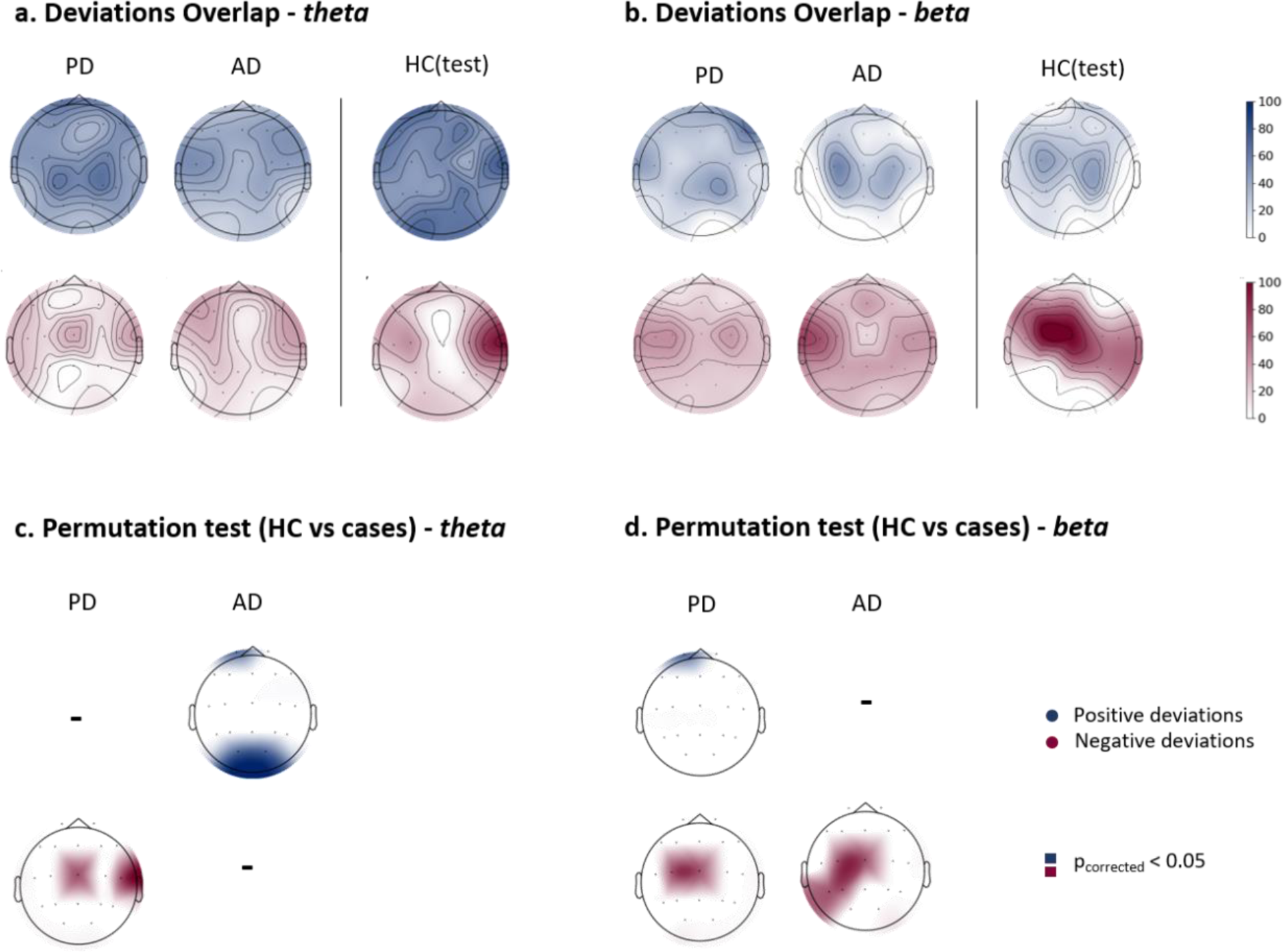
Relative power overlap maps. **(a, b)** Overlap maps of deviation scores for clinical groups and the held-out healthy control group (HC(test)), illustrating areas of common deviation in theta and beta bands respectively. (**c, d)** Significant overlap maps highlighting channels with significant differences between HC(test) and clinical groups, determined by group-based permutation tests (*p<0.05*, *FDR corrected*) in theta and beta, respectively.

We then compared the channel-wise overlap maps between PD and AD groups and the HC group using group-based permutations (Segal et al. 2023), generating significant overlap maps (Fig. 4 c,d). All results were corrected for multiple comparisons using FDR (*p* < 0.05). Our results showed that the overlap maps of both clinical groups, PD and AD, differed significantly from the HC(test) group at the delta, theta, and beta bands. In general, PD patients shared several deviating fronto-central channels for EEG power biomarker (e.g., 6 out of 19 channels) at the delta band, and 2-3 deviating central channels at the theta and beta bands respectively. As for the AD group, few channels survived the permutations and correction at the delta band, while some occipital channels at the theta band and 4 out of 19 left temporal parietal channels at the beta band were significant. Results of overlap maps and permutations test for the remaining frequency bands are presented in Supplementary Fig. S11-S13.

Regarding the EEG functional source connectivity features, the spatial overlap of extreme deviations across participants with at least one extreme deviation was notably low, not exceeding 25% across all cases and EEG frequency bands, as shown in Fig. 5. This indicates low consistency in the spatial location of extreme EEG functional source connections among individuals within the same group. Significant differences, detected through group-based permutation tests and corrected using FDR, are illustrated in Fig. 5 for the theta and beta bands and in Supplementary Fig. S14-S16 for the other frequency bands. These results illustrate the variations in the overlap maps between clinical (i.e., PD and AD) and HCy groups. Notably, only EEG source functional connections with extreme negative deviations survived the permutations and correction tests.

**Fig. 5.**
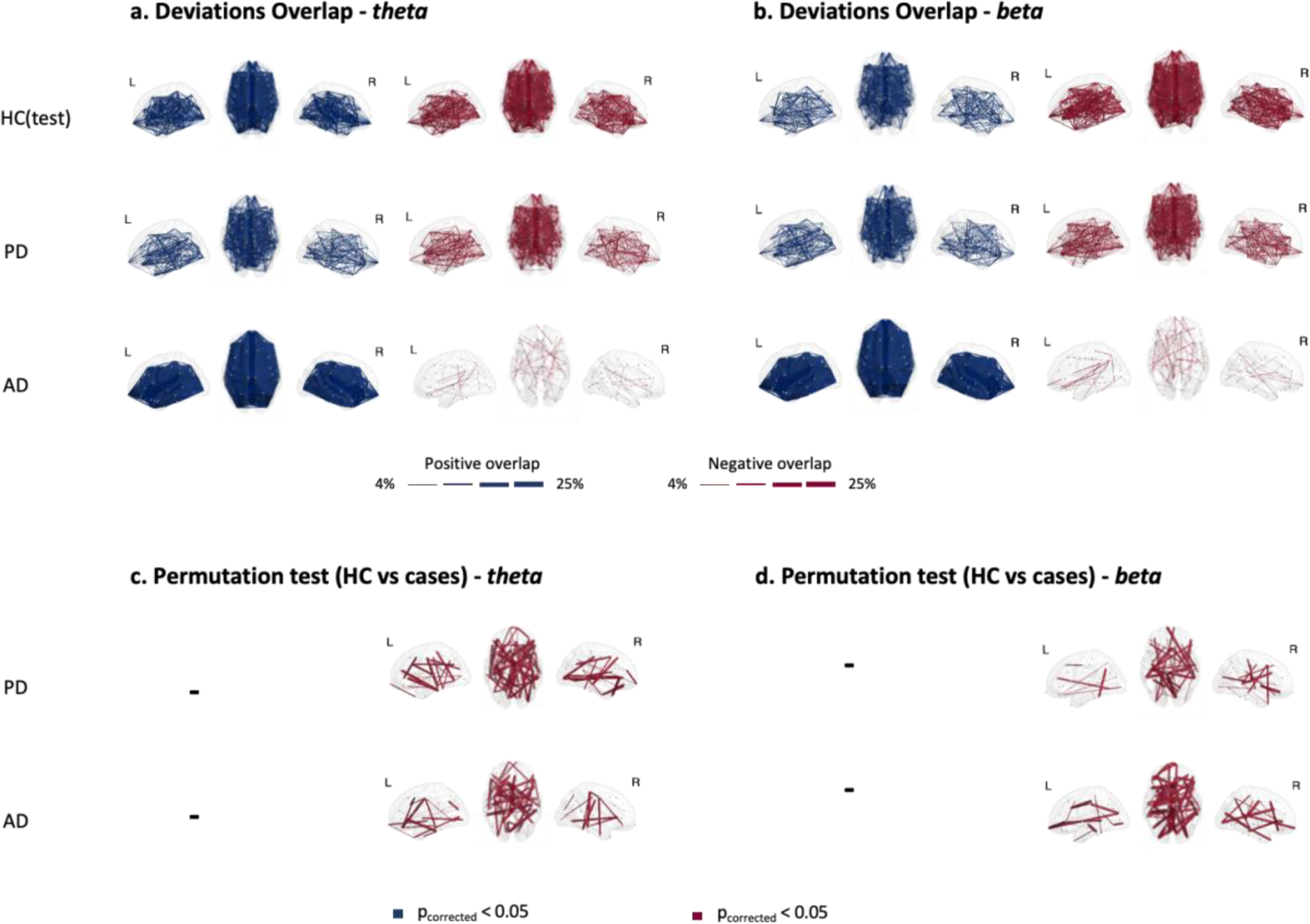
Functional connectivity overlap maps. **(a,b)** Overlap maps of positive and negative deviation scores for clinical groups and the held-out healthy control group (HC(test)) within theta and beta bands respectively, illustrating areas of common deviation among patients. **(c,d)** Significant overlap maps of functional connections showing significant differences between HC(test) and clinical groups at theta and beta bands, determined through group-based permutation tests (*p<0.05*, *FDR corrected*).

The results revealed that the majority of significantly deviated EEG source functional connections belonged to the default mode network (DMN) network, particularly at the delta (PD: 39%, AD: 44%), theta (PD: 40%, AD: 35%) and alpha (PD: 42%, AD: 44%) bands for both clinical groups. Detailed proportions across resting-state cortical source networks for each clinical group and frequency band can be found in Supplementary Table S8.

### Normative model-derived scores as patient-specific markers for clinical assessment

Given the above demonstrated high within-group heterogeneity of the EEG biomarkers used, it is crucial to take into account the participant-specific deviations in any further analysis. Thus, we used the patient-specific deviations to compute a new metric called EDI (extreme deviations index) defined as the averaged EEG source connectivity of the extremely deviated solutions. This metric was then correlated (Spearman’s correlation) with the patients’ clinical assessments. For the PD patients, we used their UPDRS scores as a clinical assessment measure. UPDRS scores were available for 86 patients. Similarly, we used the MMSE scores to assess the global cognitive function in the AD patients, with MMSE scores available for a total of 146 AD patients. In addition, we examined the correlation between the EDI and the general cognitive status of the PD patients using the available MMSE scores from 182 PD patients. We found a significant correlation at the delta band for both PD patients (*⍴*=0.24, *p*=0.0254) and AD patients (*⍴*=-0.26, *p*=0.0107), as depicted in Fig. 6. These findings represent a preliminary step toward developing EEG-based patient-specific markers that can be used for objectively quantifying personalized treatment modalities. More details on the AD subjects showing significant *p* with MMSE scores in other frequency bands can be found in supplementary Fig. S17-S18.

**Fig. 6.**
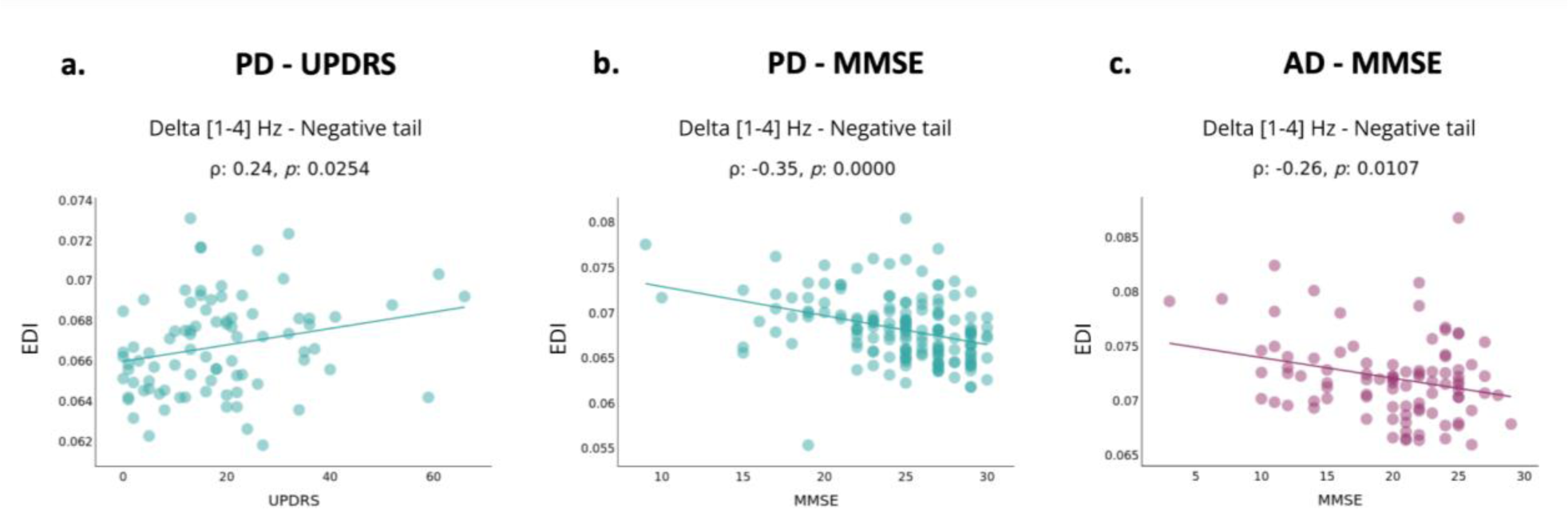
NM-derived scores for clinical assessments. (a) Correlation between PD subjects’ ‘EDI’ and UPDRS in the delta band for negative extreme deviations. (b) Correlation between PD subjects’ ‘EDI’ and MMSE in the delta band for negative extreme deviations. (c) Correlation between AD subjects’ ‘EDI’ and clinical assessment scores (MMSE) in the delta band for negative extreme deviations.

## Discussion

In this study, we aimed to achieve three key contributions. First, we mapped the aging trajectories of the EEG spectral power at scalp electrodes and EEG functional connectivity at cortical source pairs in a HC cohort (>40 yo) as popular neurophysiological biomarkers of interest. Second, we demonstrated the electrophysiological heterogeneity among PD and AD patients, challenging the prevailing reliance on the traditional case-control approach, and emphasizing the importance of acknowledging individual variability. Last, the identification of patient-specific markers derived from normative models showed an association with clinical assessments, which constitutes a preliminary step in developing EEG patient-specific biomarkers for those neurodegenerative diseases with potential applications to other neurodegenerative diseases.

### Electrophysiological normative aging maps

We used a normative modeling approach to map the trajectories of EEG biomarkers (features) of interest throughout the >40yo lifespan, providing a reference model to which individual cases can be compared. As mentioned above, we selected EEG relative spectral power and functional source connectome across standard frequency bands from delta to gamma as response variables for the normative models due to their notable prevalence in the EEG literature (Chiarion et al. 2023; Zhang et al. 2023) and the substantial body of research reporting age-related alterations in these features during the process of normal and pathological aging (Benwell et al. 2020; Harada, Natelson Love, and Triebel 2013; Javaid, Kumarnsit, and Chatpun 2022; Meghdadi et al. 2021).

For both EEG features and across all frequency bands, we observed continuous trajectories across age with relatively low variability in averaged EEG functional source connectivity features compared to spectral features. These developmental trajectories suggest that the healthy human brain undergoes gradual development, maintaining a state of stability as individuals age beyond 40 yo. The slow developmental trajectory observed in our study may be attributed to the brain’s adaptive capacity, accumulated knowledge, and lifelong experiences that shape brain function over time (Harada, Natelson Love, and Triebel 2013; Hedden and Gabrieli 2004; Oschwald et al. 2019; Salthouse 2010).

After constructing the typical aging trajectories from the HC cohort, we sought to elucidate abnormal changes in the electrophysiological aging maps in neurodegenerative diseases. We focused on characterizing the extent to which patient metrics deviate from established norms, thereby providing preliminary insights into the clinical utility of our electrophysiological normative models. Emerging evidence increasingly suggests abnormal alterations in EEG spectral power (Al-Qazzaz et al. 2014; Benwell et al. 2020; Chaturvedi et al. 2017; Lejko et al. 2020; Shirahige et al. 2020; McMackin et al. 2019) and functional connectivity (Briels et al. 2020; Lejko et al. 2020; McMackin et al. 2019) associated with AD and PD. Significant deviations in EEG functional source connectivity within default mode network (DMN) and dorsal attention network (DAN) were observed in both clinical groups across frequency bands. Based on the comprehensible review by (Filippi et al. 2023), research work has documented disruptions within these networks in the context of neurodegenerative diseases, reflecting their pivotal roles in cognitive processing and their susceptibility to age-related pathological changes.

### Heterogeneity in EEG spectral and connectivity features

Despite the considerable body of research on the induced alterations in EEG features associated with AD and PD, findings across studies lack consistency. Specifically, studies focusing on AD patients have reported examples of inconsistencies in resting-state EEG functional source connectivity literature (Briels et al., 2020). This variability may be attributed to the lack of a standardized methodology for EEG acquisition and functional analysis, as well as the reliance on dataset samples that may compromise the results’ generalizability. We think that a primary contributor to this inconsistency arises from the inherent heterogeneity among patient populations affected by PD and AD (and other neurodegenerative diseases), which introduces confounding factors often inadequately addressed in group-level analyses. For instance, despite the general trend of EEG power density shift from higher to lower frequency bands observed in neurodegenerative disease groups compared to HC controls, as outlined in recent systematic reviews (Lejko et al. 2020; Shirahige et al. 2020), numerous studies have revealed individual heterogeneity among patients with aging pathologies (Chen et al. 2024; Duara and Barker 2022; Robinson et al. 2023; Vogel et al. 2023; Wüllner et al. 2023). The main contributors to this substantial heterogeneity are phenotypic diversity, reflecting disease subtypes, and temporal variability, including various disease stages, sex, education attainment, and several endogenous and environmental disease risk factors (Hampel et al. 2018; Young et al. 2018). In our study, the identified heterogeneity in aging EEG functional patterns, characterized by an overlap below 25% for the deviated EEG functional source connectivity measures, aligns with recent findings from investigations that utilized structural MRI and normative modeling in aging cohorts. Indeed, it was shown that spatial patterns of cortical thickness were heterogeneous among AD patients, with overlapping outliers at the regional level not exceeding 50% within the AD group (Verdi et al. 2023). We believe that the demonstrated heterogeneity poses a significant barrier to the development of EEG-based biomarkers. The challenge associated with the group-level analysis manifests across multiple levels. At the diagnostic level, patients are typically categorized into distinct groups, assuming uniformity within each group. Similarly, treatment approaches often adopt a general ‘*one-size-fits-all*’ strategy, overlooking individual differences. To tackle this issue effectively, the development of patient-specific electrophysiological biomarkers is imperative.

### Beyond heterogeneity mapping

It is crucial to recognize that the differentiation between neurodegenerative diseases, particularly during early and middle stages, poses a significant challenge in clinical practice due to overlapping clinical symptoms across different conditions (Armstrong, Lantos, and Cairns 2005). In particular, individuals with mild cognitive impairment (MCI) may share common clinical and neurophysiological profiles among aging pathologies, adding complexity to capturing the nuanced variations between conditions (Jongsiriyanyong and Limpawattana 2018). Our work addresses this challenge by generating electrophysiological aging trajectories and leveraging normative models to derive patient-specific deviation scores, which may offer a refined clinical approach compared to traditional methods relying solely on raw features. In the framework of Precision Medicine, the patient’s assessment is aimed at not only providing a clinical diagnosis by proper in-vivo biofluid or neuroimaging techniques measuring disease-specific neuropathological species in the brain of PD or AD patients (e.g., amyloid-beta and tau in AD patients), but also to use reliable and valid biomarkers accounting for neurobiological, neuroanatomical, and neurophysiological underpinnings of the individual clinical manifestations along the disease course. Along this line, here we proposed normative models and neurophysiological measures of deviance from resting-state EEG rhythms reflecting specific alterations in the regulation of quiet vigilance in PD and AD patients that are quite relevant to the quality of life of patients (e.g., watching TV programs, reading books and newspapers, having quiet social conversation, etc.) (Babiloni 2022).

It is noteworthy to mention that our clinical dataset comprises AD and PD patients with MCI. For example, Dataset 7 includes 38 PD patients with MCI as well as 13 AD patients with MCI. However, the subtype distribution of patients in other datasets is not explicitly stated, potentially including cases of dementia and MCI. As part of our future endeavors, we aim to elucidate patient data regarding subtypes further and use deviation scores derived from the normative models to distinguish between subtypes within each disease, motivated by the work of Young et al., who sought to develop a machine learning tool that enables the discovery of disease subtypes and stages for precise medicine in neurodegenerative disorders (Young et al. 2018).

Furthermore, a primary objective of combining EEG and normative modeling is to establish a patient-specific marker with clinical utility. Assessing whether extreme deviations in functional metrics are associated with symptom severity or cognition is crucial. In this study, we examined the clinical correlates of extremely deviated features and demonstrated significant associations between individual-specific deviations and clinical assessments. Thus, normative models hold promise for the development of personalized electrophysiological approaches.

### Limitations

Several challenges warrant further consideration. First, while our sample size is substantial for EEG studies (n∼933), it does not reach the scale often seen in the most important international MRI and fMRI initiatives, and it may not be fully representative of the general population. Notably, in this study, EEG data collected from HC participants primarily fall within the age range (60-70 years), with a notable lack of EEG data above 72 years (see Fig. 1). Further efforts should incorporate additional neuroimaging cohorts to achieve a more balanced representation of individuals across diverse age ranges. We anticipate that EEG-based aging maps established herein will serve as a dynamic resource, with ongoing updates to the electrophysiological aging model as more resource data sets become available. We will specifically focus on integrating longitudinal datasets for a precise characterization of aging developmental trajectories.

Our data was gathered from 14 distinct spontaneous EEG studies, each using different EEG systems with a varying spatial resolution (19, 32, 64, 128, 256 channels) and protocol and instructions to participants not harmonized. To standardize our analysis for scalp spectral analysis of EEG activity, we mapped all systems to a common low spatial resolution (19 channels). Yet, this approach may overlook some relevant information that could be derived from higher spatial resolution setups. Moreover, sample sizes varied across datasets. However, to address this issue and identify potential bias arising from individual datasets, we ran a leave-one-study-out analysis. Results showed that no specific dataset significantly influenced NM trajectories (See Supplementary Fig. S5, S6). Another methodological limitation was the use of fixed EEG frequency bands for delta, theta, and alpha despite the recommendations on the spectral analysis of resting state EEG rhythms by expert panels of the International Federation of Clinical Neurophysiology and The Alzheimer’s Association International Society to Advance Alzheimer’s Research and Treatment (Babiloni et al. 2020; 2021; 2021). These recommendations are based on the evidence that compared to HC persons, AD and PD patients have a remarkable frequency slowing of EEG alpha rhythms (even more pronounced in PD than AD patients), suggesting the use of individual alpha frequency peaks as a landmark to define delta, theta, and alpha frequency bands. In the present study, we preferred to use standard frequency bands for delta, theta, and alpha rhythms to allow a more direct comparability with the large majority of the EEG literature on PD and AD.

Another challenge of the study arises from the inherent heterogeneity within patient groups, with individuals exhibiting different stages of PD and AD diagnosis. For instance, many datasets lack sufficient information to confirm whether PD or AD patients were at MCI or dementia stages. Therefore, we focused on examining disease-related electrophysiological normative model metrics, regardless of the disease stage. Additionally, a limitation of this study is the potential influence of medications on EEG results in the patient cohort. While we aim to uncover neurobiological patterns, it is important to acknowledge that medication effects could introduce part of the variability shown in the present study. Addressing individual variations and medication interactions was challenging due to incomplete medication profiles in some cases. Future studies could be improved by including detailed records of disease stages and pharmacological histories. In addition, it is recommended for future studies to consider the diversity of environmental factors as covariates in normative modeling, particularly given that data is collected from multiple sites with different socio-economic statuses and education levels, which may influence brain aging trajectories.

## Conclusion

This study utilized popular resting-state EEG biomarkers and normative modeling to delineate the typical aging trajectories of EEG spectral power and cortical functional source connectivity features and explore the electrophysiological heterogeneity in PD and AD patients. Our findings revealed significant variability in those EEG features among patients, with deviations from normative trajectories correlating with the clinical severity. These findings emphasize the emergent need for patient-level inferences to enhance the accuracy of the neurophysiological assessment in PD and AD patients and inform more personalized treatment strategies. Further research and clinical validation will be necessary to realize these potential benefits fully.

## Materials and Methods

### Datasets

Our cohort consisted of 933 individuals, subdivided into a group of healthy controls (N=400 in the training set, N=99 in the held-out testing set) and a group of 434 participants clinically diagnosed with neurodegenerative disorders, including PD (N=237), and AD (N=197). All participants were aged above 40 years old (mean = 64.95 ± 10.28; 48% M) and underwent resting-state EEG recordings with their eyes closed. For subjects with multiple sessions/runs, only the first session/run was included. The data were aggregated from 14 distinct datasets, with each being approved by its local ethics committee. Please refer to Supplementary Table S1 for a detailed and comprehensive overview of the datasets used in this study, and Fig. 1 for age distribution across groups, sex, and sites.

### Data Preprocessing

The EEG preprocessing and artifact removal pipeline employed a multi-stage and automated algorithm, supported by visual inspection. Initially, EEG signals underwent bandpass filtering (1-100 Hz). All data signals were downsampled to a common frequency (200 Hz), and a notch filter was applied to target the dataset-specific line frequency. Bad EEG channels were identified using the *pyprep* algorithm, which employs a *RANSAC*-based approach, and interpolated based on neighboring electrode data (Bigdely-Shamlo et al. 2015). *RANSAC* selects a small group of EEG channels, estimates a model based on these channels, and then identifies potential outliers or bad channels. Next, re-referencing was applied using the common average reference method to minimize noise across electrodes. Eye blink artifacts were identified and rejected using Independent Component Analysis (ICA) the *IClabel* algorithm (Pion-Tonachini, Kreutz-Delgado, and Makeig 2019). A second bandpass filter (1-45 Hz) further refines the data. Then, EEG signals were segmented into 10-second epochs as a trade-off between the needed length for computing the connectivity matrices and the available segment length per site, and the *Autoreject* toolbox (Jas et al. 2017) was used to detect and clean or reject bad epochs. All EEG datasets underwent the same preprocessing steps described here, except for dataset 3 (*BASEL*), which was already preprocessed as detailed in (Yassine et al. 2022).

### Features Extraction

#### Scalp-level Spectral features

The normative model establishes the relationship between a response variable and one or more covariates. In this study, we initially focused on the spectral features of the EEG signal as the designated response variable. This choice was motivated by the extensive literature highlighting the changes in EEG power associated with neurodegenerative diseases (Chaturvedi et al. 2019; Lejko et al. 2020; Shirahige et al. 2020; Al-Qazzaz et al. 2014). The power spectrum density (PSD) for each epoch and each channel is computed using Welch’s method (1-second *Hann* window with a 50% overlap, and a spectral resolution of 0.5 Hz). PSDs are then averaged across all epochs within a subject. Relative power in specific frequency bands (delta [1-4 Hz], theta [4-8 Hz], alpha [8-13 Hz], beta [13-30 Hz], gamma [30-45 Hz]) is computed by dividing the absolute power within each narrow band by the total power of the broader band [1-45 Hz]. Since the number of channels was not consistent across all datasets, we downsampled the electrode configurations to a common 10-20 montage, consisting of 19 channels.

#### Source-level Functional connectivity features

We computed EEG-based functional networks using the EEG source connectivity method (Hassan and Wendling 2018). Cortical sources were estimated using the exact low-resolution brain electromagnetic tomography (eLORETA) (Pascual-Marqui 2007). The noise covariance matrix was set to an identity matrix and the regularization parameter was fixed at λ=0.1. Age-specific head models of the brain, skull, and scalp layers were built using an MRI template of elderly individuals aged 65-69 years (Fillmore, Phillips-Meek, and Richards 2015), employing the Boundary Element Method (BEM) from the MNE Python package. The forward and inverse models were solved within a source space of 4098 sources per hemisphere, with approximately 5 mm spacing. We then downsampled the source space to 68 representative sources by averaging the sources within each region defined by the Desikan-Killiany atlas (Desikan et al. 2006). Subsequently, we computed the functional connectivity between pairwise regions of interest, using the amplitude envelope correlation (AEC) method, defined as the Pearson correlation between signals’ envelopes derived from the Hilbert transform (Brookes et al. 2011; Hipp et al. 2012). To mitigate zero-lag signal overlaps caused by spatial leakage, we applied a pairwise orthogonalization approach before computing connectivity (Brookes, Woolrich, and Barnes 2012).

### Normative Modeling

Normative Modeling (NM) aims to establish a normative relationship between a response variable (behavioral, demographic, or clinical variables) and at least one covariate (a quantitative biological measure, e.g. age or sex). To estimate the normative age-related curves for EEG spectral power and functional connectivity (as response variables), we implemented the Generalized Additive Models for Location, Scale, and Shape (GAMLSS) (Stasinopoulos and Rigby 2008) using the *gamlss* package. We started by identifying the optimal data distribution and best-fitting parameters and covariates. Utilizing these specific GAMLSS models, we obtained nonlinear normative trajectories for each feature and at each frequency band. The model performance was assessed by the model convergence, residuals, and Q-Q (quantile-quantile) plots. The model sensitivity was analyzed with a leave-one-study-out (LOSO) analysis. Leveraging these population-level normative trajectories, we established benchmarks for each subject using individualized deviation scores.

### GAMLSS framework

GAMLSS are semi-parametric regression models offering a flexible framework for capturing complex relationships (Rigby and Stasinopoulos 2005). They assume a specific distribution for the response variable, with parameters linked to a set of explanatory variables through linear or nonlinear predictor functions. The mathematical formulation of GAMLSS is as follows:

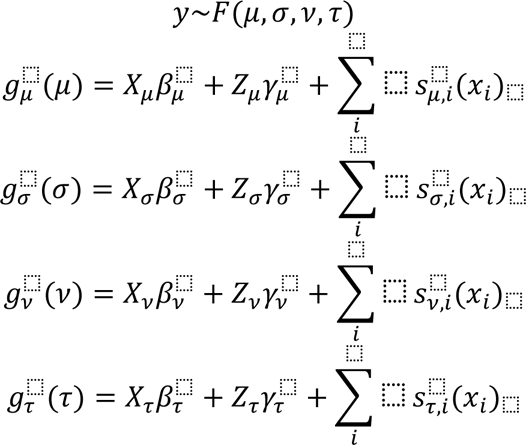

The response variable *y* follows a distribution *F* defined by parameters (*μ*, *σ*, *v*, *τ*). Each parameter is linked to explanatory variables via the link function *g*(), where *β* represents the fixed effect term and *X* is its design matrix. *γ* accounts for the random effects, and Z is its design matrix. *s* denotes the non-parametric smoothing function (Bethlehem et al. 2022; Rigby and Stasinopoulos 2005). In this study, our response variable is an EEG-derived feature, and age serves as the main covariate. The inclusion of other covariates such as sex and data collection sites is detailed in subsequent sections. Following Bethlehem et al., we used fractional polynomials as a smoothing function to accommodate nonlinearity while maintaining model stability (Bethlehem et al. 2022).

### Model distribution

The GAMLSS framework provides an extensive range of distribution families. Here, we used an empirical approach to determine the most suitable distribution, by training models across all considered distribution families (with 3 or more moments, continuous/mixed), and comparing them using the Bayesian Information Criterion (BIC). The distribution with the lowest BIC score was selected. This process was systematically applied to the two features under study. The distributions that best fit the averaged spectral power and connectivity values are reported in supplementary Tables S4, and S5, respectively. In addition, we determined the optimal number of polynomials for the age covariate and whether to include it in parameters beyond μ by comparing BIC scores across various models.

### Model covariates

Model covariates beyond age, including sex and site (considered both as a fixed effect and a random effect), are empirically selected. Each covariate is sequentially incorporated into the parameter formulas, and the resulting models are compared based on their BIC scores. The model with the lowest BIC score is chosen, determining whether the covariates are retained in the final model. The final models for spectral and connectivity features are reported in supplementary Tables S4 and S5. See Fig. S1, S2, and Tables S2, S3 for an exploratory analysis of the differences in EEG feature distributions between male and female participants within each group.

### Model performance

To evaluate the performance of our models, we examined the normalized quantile residuals. Visual inspection of the residual plots depicted in Supplementary Fig. S3, S4 suggests that our models exhibit adequate fit and quality. Specifically, the residuals plotted against the fitted values of *μ* and the index were evenly scattered around the horizontal line at 0. The kernel density estimation of the residuals displayed an approximate normal distribution, and the normal quantile-quantile (Q-Q) plots showed an approximately linear trend with an intercept of 0 and a slope of 1.

### Model sensitivity

To validate the robustness of the model, we conducted a series of leave-one-study-out analyses. Specifically, we systematically excluded one dataset from the primary datasets, refitted the GAMLSS models, evaluated all model parameters, and then extracted developmental trajectories. We then compared these alternative trajectories to those derived from the 14 datasets for each feature and frequency band. Our findings showed remarkable consistency, with a very high correlation between the trajectories derived from the primary full dataset and those from the subsets (all r > 0.88 for spectral features, all r > 0.90 for FC features), even when large datasets were excluded (Supplementary Fig. S5, S6).

### Deviation maps

After parameters selection and validation, we trained GAMLSS models for each channel/connection across all frequency bands using the healthy control training group HC(train) (80% of the healthy sample). Subsequently, we projected the features data (i.e., relative power and functional connectivity values), of our clinical groups (PD, and AD) and the held-out healthy control testing group HC(test) (remaining 20% of the healthy sample), onto the corresponding models. This process enables us to generate the individual centiles and calculate the deviation scores (z-scores) using the quantile randomized residuals approach (Dunn and Smyth 1996), for each channel/connection and each subject, resulting in an individual deviation map per subject.

### Overlap maps

An extreme deviation is defined as |z-score| > 2. Consequently, we derived positive and negative extreme deviation maps for z-scores > 2 and < -2, respectively. We then computed the number of subjects with at least one extreme deviation, as well as the number of extreme deviations per subject. Additionally, for each channel and location, we calculated the percentage of subjects showing extreme deviations at this specific location among those with at least one extreme deviation, which resulted in group-specific overlap maps.

### Significant overlap maps

We used group-based permutation tests to assess group differences in channel/connectivity-level overlap maps (Segal et al. 2023). This involved shuffling case and control labels of individual-specific deviation maps. During each iteration, group labels were permuted, resulting in a new grouping of extreme deviation maps for each subject based on the shuffled labels. Subsequently, new overlap maps were computed for both HC(test) and clinical groups. By subtracting the surrogate HC(test) overlap map from the surrogate clinical group’s overlap map, an overlap difference map for each disorder was derived. This procedure was repeated 5,000 times to establish an empirical distribution of overlap difference maps under the null hypothesis of random group assignment. For each channel/connection, *p*-values were obtained as the proportion of null values that exceeded the observed difference. Statistically significant effects were identified using two-tailed FDR correction (*p<0.05)*.

## Data availability

Datasets 1, 2, 4, 11, 12, 13, and 14 are openly accessible on the OpenNeuro, OSF, and Zenodo platforms. The corresponding download links can be found in Supplementary Table S1. Access to other datasets can be made available upon request.

## Code availability

Codes are available at https://github.com/MINDIG-1/NM-neurodeg. We used *gamlss* package in R (Stasinopoulos and Rigby 2008) for statistical modeling, *MNE-python* package (https://mne.tools/stable/index.html) for EEG signal processing, and BrainNet Viewer (https://www.nitrc.org/projects/bnv/) (Xia, Wang, and He 2013) for networks visualization.

## Acknowledgments

This work was fully funded by MINDIG as a part of its R&D activity. We would like to thank all the researchers who shared their data in open access and all the participants who approved the use of their data in research. This work was supported by the ‘Region Bretagne’, Inno R&D project n 23001155 and Rennes Metropole (AICE project).

## Author contributions

J.T., S.A., AE. and M.H. conceived the study and wrote the manuscript, with valuable revision from all authors. Results were interpreted by J.T., S.A., A.M., A.E, C.B. and M.H., with contributions from all authors. A.M. M.V., B.G., G.Y, U.G, P.F., V.P. provided datasets. A.K. and A.E. helped in the data preprocessing and analysis part.

## Competing interests

N/A

## Footnotes

1 https://population.un.org/wpp/Graphs/DemographicProfiles/Line/900

2 https://www.alzint.org/news-events/news/new-data-predicts-the-number-of-people-living-with-alzheimers-disease-to-triple-by-2050/

3 https://news.umiamihealth.org/en/parkinsons-disease-where-does-it-start-and-why-does-it-matter/

## Notes

### Competing Interest Statement

The authors have declared no competing interest.

