## Supplementary for "Characterizing the heterogeneity of neurodegenerative diseases through EEG normative modeling"

### Methods

#### Datasets

Our cohort consisted of 933 individuals, subdivided into a group of healthy controls (n=400 in the training set, n=99 in the held-out testing set) and a group of 434 participants clinically diagnosed with neurodegenerative disorders, including PD (n=237), and AD (n=197). The data were aggregated from 14 distinct studies, each detailed in Table S1.

**Table. S1 | Demographic and EEG system details of the control and clinical cohorts used in this study across 14 sites.**

Supplementary_Table_S1 (Excel File)

### Exploratory Data Analysis

We performed a Mann-Whitney U test to check for any significant differences in the EEG spectral power and functional connectivity distribution between male and female participants within each group. The effect sizes of these differences were quantified using Cohen's d.

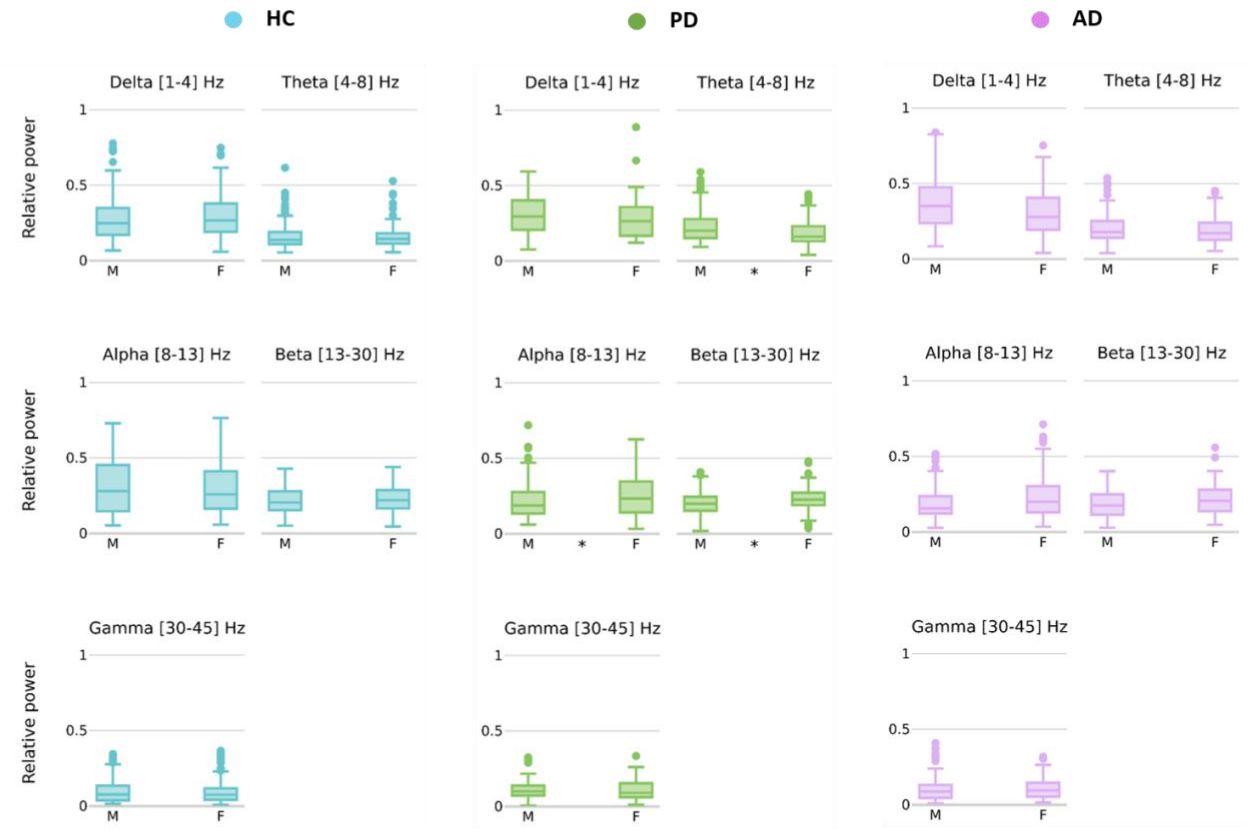


**Fig. S1 | Sex differences in the averaged relative power of EEG frequency bands in Healthy Controls (HC), Parkinson's Disease (PD) and Alzheimer's Disease (AD) groups.** (*) denote statistically significant differences between sexes within each frequency band.

**Table. S2 | Sex differences in the averaged relative power of EEG frequency bands: *P-value (Cohen's d).***

| | Delta [1-4] Hz | Theta [4-8] Hz | Alpha [8-13] Hz | Beta [13-30] Hz | Gamma [30-45] Hz |
|---|---|---|---|---|---|
| **HC** | 0.06 (-0.15) | 0.75 (0.04) | 0.63 (0.07) | 0.16 (-0.11) | 0.39 (0.22) |
| **PD** | 0.09 (0.18) | 0.00 (0.46) | 0.02 (-0.35) | 0.01 (-0.33) | 0.70 (0.03) |
| **AD** | 0.06 (0.29) | 0.42 (0.18) | 0.06 (-0.32) | 0.11 (-0.25) | 0.46 (0.04) |

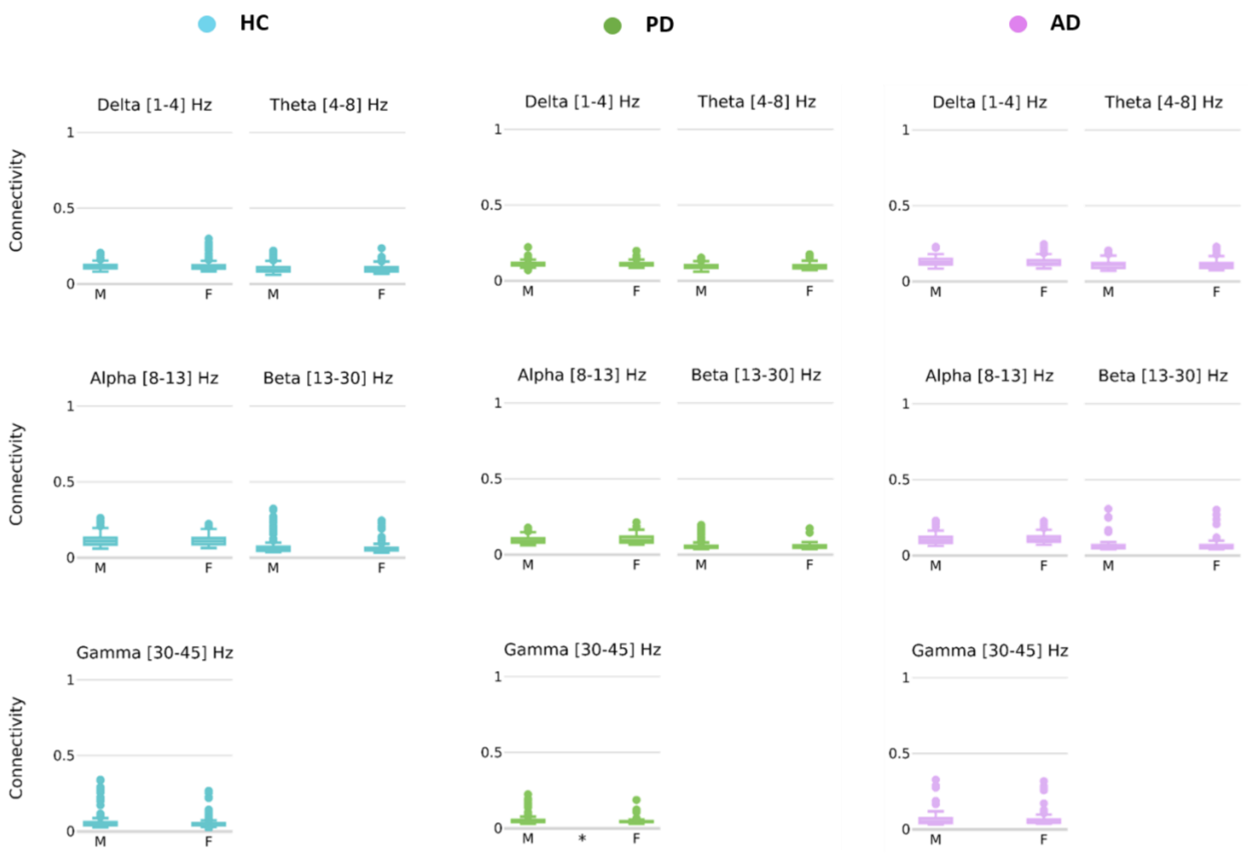


**Fig. S2 | Sex differences in the averaged functional connectivity of EEG frequency bands in Healthy Controls (HC), Parkinson's Disease (PD) and Alzheimer's Disease (AD) groups.** (*) denote statistically significant differences between sexes within each frequency band.

**Table. S3 | Sex differences in the averaged functional connectivity of EEG frequency bands: *P-value (Cohen's d)*.**

| | Delta [1-4] Hz | Theta [4-8] Hz | Alpha [8-13] Hz | Beta [13-30] Hz | Gamma [30-45] Hz |
|---|---|---|---|---|---|
| **HC** | 0.27 (-0.05) | 0.81 (0.05) | 0.84 (0.06) | 0.37 (0.25) | 0.11 (0.26) |
| **PD** | 0.22 (0.12) | 0.45 (-0.06) | 0.11 (-0.28) | 0.61 (0.08) | 0.00 (0.28) |
| **AD** | 0.07 (0.17) | 0.78 (-0.01) | 0.27 (-0.12) | 0.96 (0.11) | 0.81 (0.11) |

### Normative Modeling

#### Model distribution and covariates

**Table. S4 | Model equations and family distribution for spectral features models across all frequency bands**

| Frequency band | Delta | Theta | Alpha | Beta | Gamma |
|---|---|---|---|---|---|
| **mu** | y ~ fp (age, 1) | y ~ fp (age, 1) | y ~ fp (age, 1) | y ~ fp (age, 1) | y ~ fp (age, 1) |
| **sigma** | ~ 1 | ~ 1 | ~ 1 | ~ 1 | ~ 1 |
| **Distribution Family** | SEP4 | GG | SEP4 | GG | GG |

**Table. S5 | Model equations and family distribution for connectivity features models across all frequency bands**

| Frequency band | Delta | Theta | Alpha | Beta | Gamma |
|---|---|---|---|---|---|
| **mu** | y ~ fp (age, 1) + factor (sex) | y ~ fp (age, 1) | y ~ fp (age, 1) | y ~ fp (age, 1) | y ~ fp (age, 1) |
| **sigma** | ~ fp (age, 1) | ~ fp (age, 1) | ~ fp (age, 1) | ~ fp (age, 1) | ~ fp (age, 1) |
| **nu** | ~ 1 | ~ 1 | ~ 1 | ~ 1 | ~ 1 |
| **Distribution Family** | ST3 | SEP3 | SEP4 | ST3 | ST1 |

#### Model performance

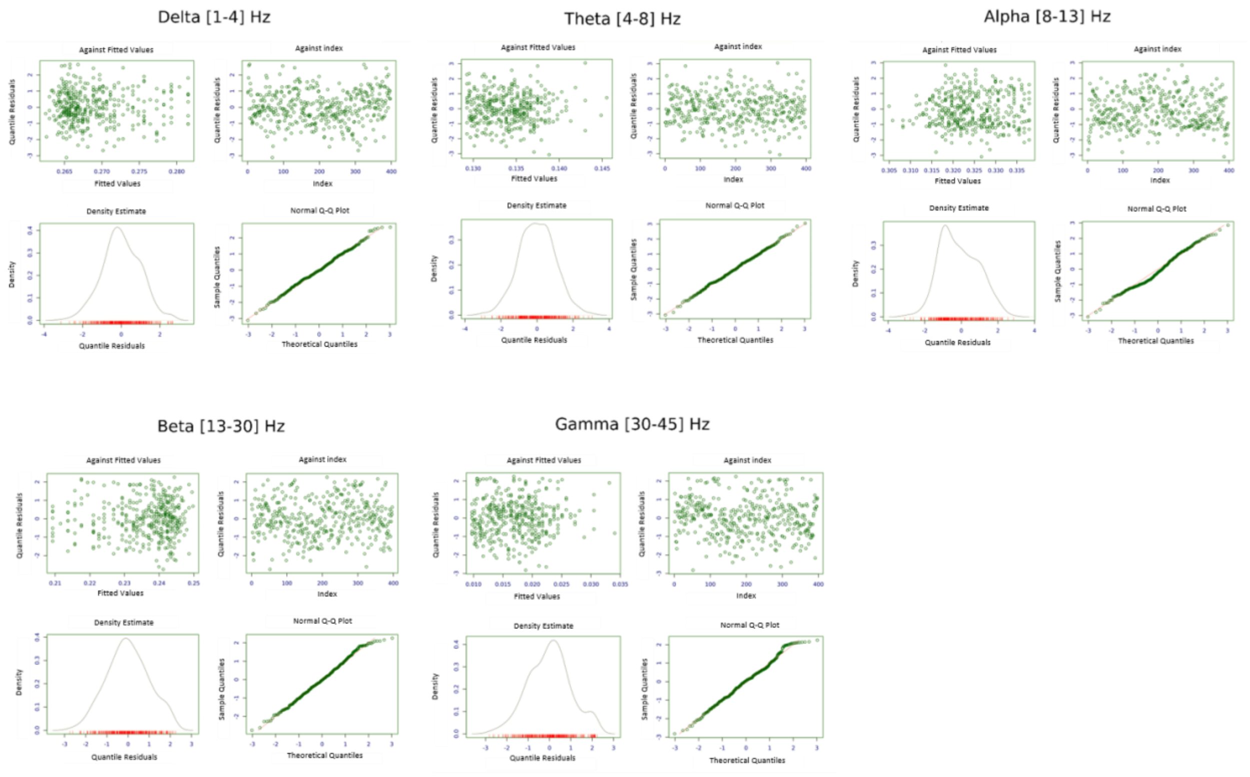


**Fig. S3 | Diagnostic residual plots for assessing spectral model fit:** residuals *vs* fitted values, index, density estimate, and normal Q-Q plot.

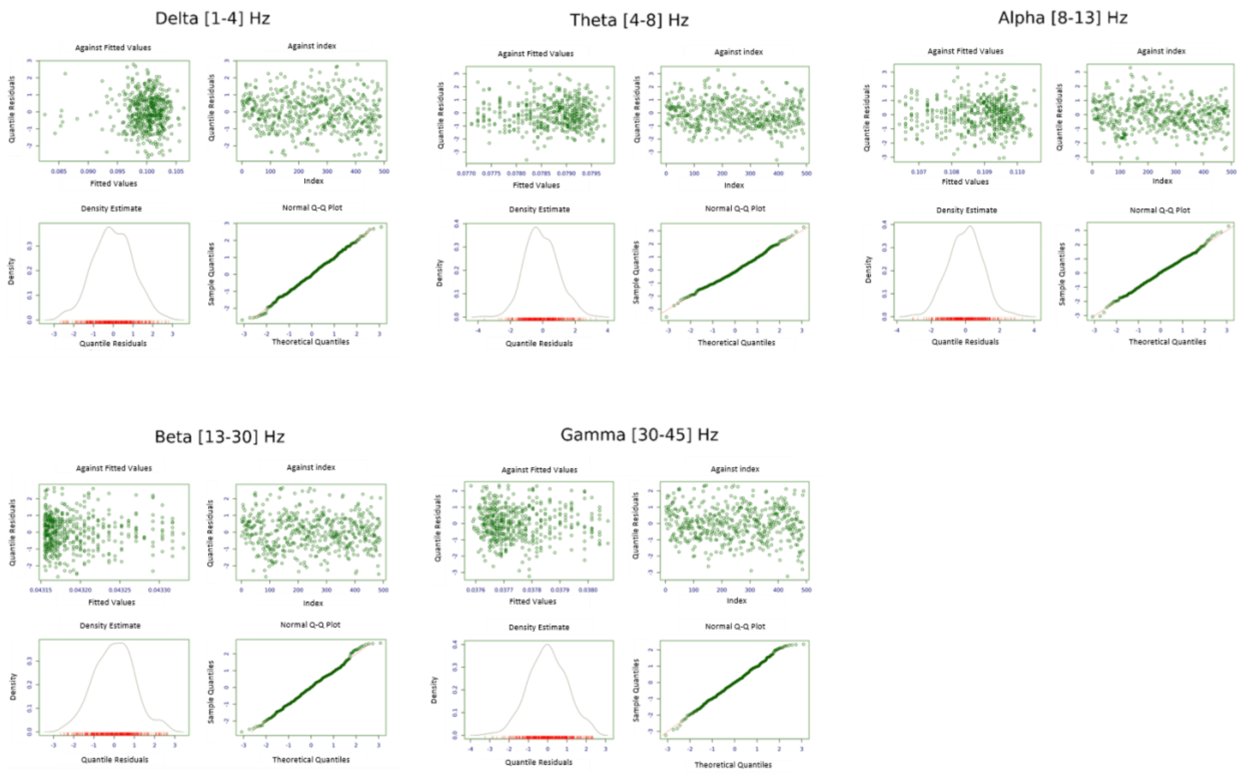


**Fig. S4 | Diagnostic residual plots for assessing connectivity model fit:** residuals *vs* fitted values, index, density estimate, and normal Q-Q plot.

#### Model Sensitivity

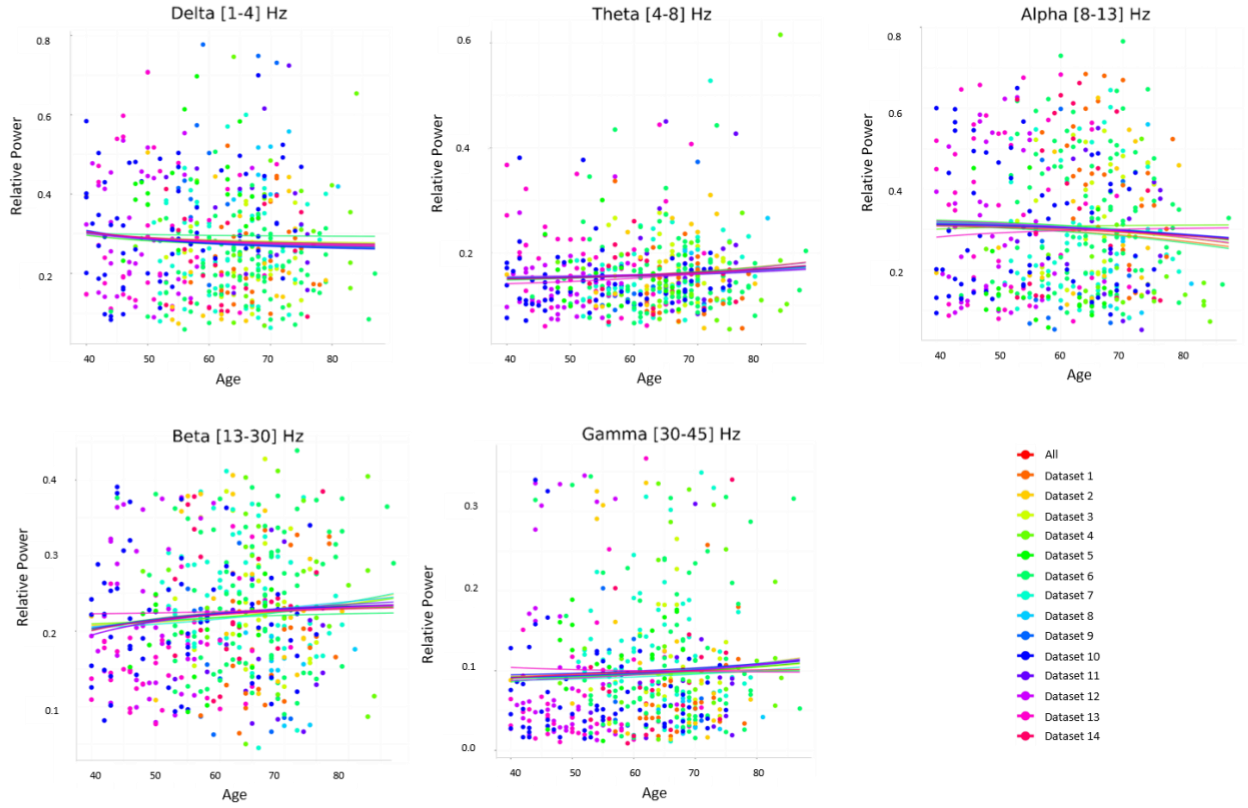


**Fig. S5 | Leave One Study Out (LOSO) analysis for assessing spectral model sensitivity on datasets:** normative model trajectories across age excluding one dataset per iteration, compared to full dataset trajectories (red) across all frequency bands.

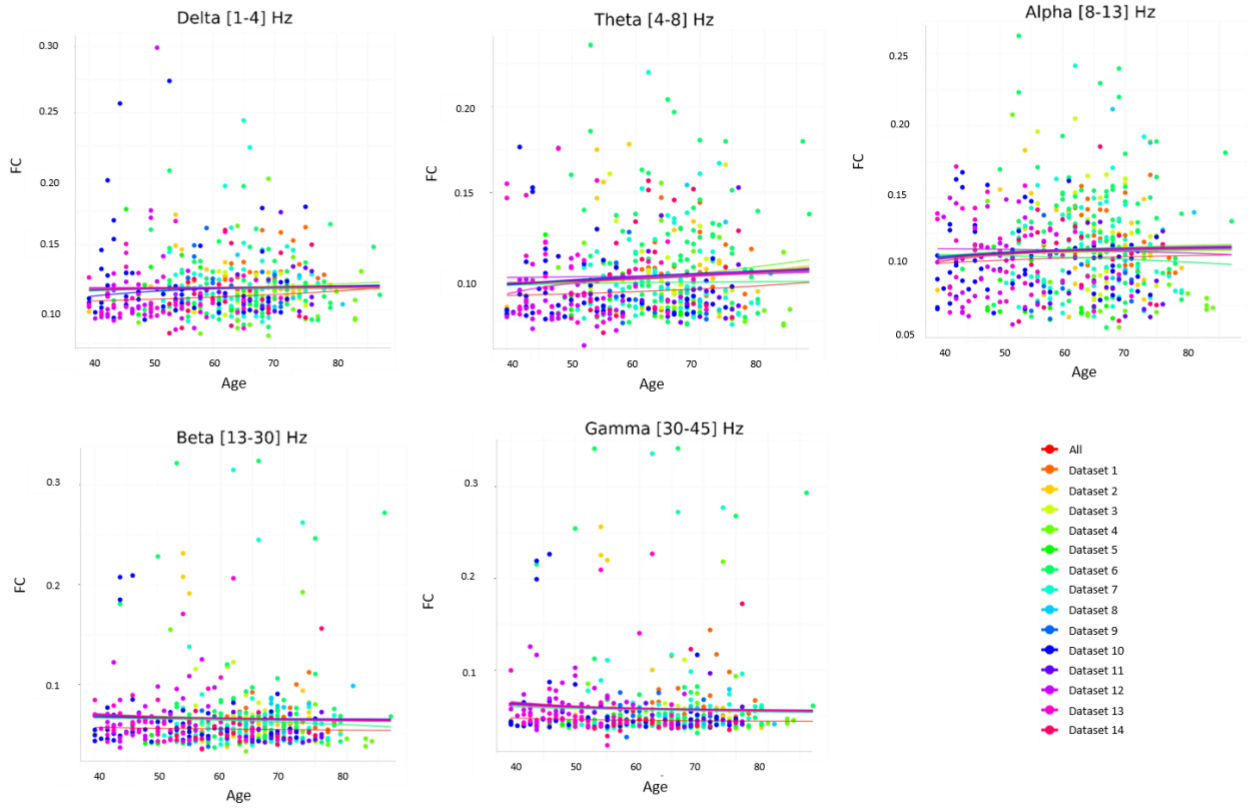


**Fig. S6 | Leave One Study Out (LOSO) analysis for assessing FC model sensitivity on datasets:** normative model trajectories across age excluding one dataset per iteration, compared to full dataset trajectories (red) across all frequency bands.

### Results

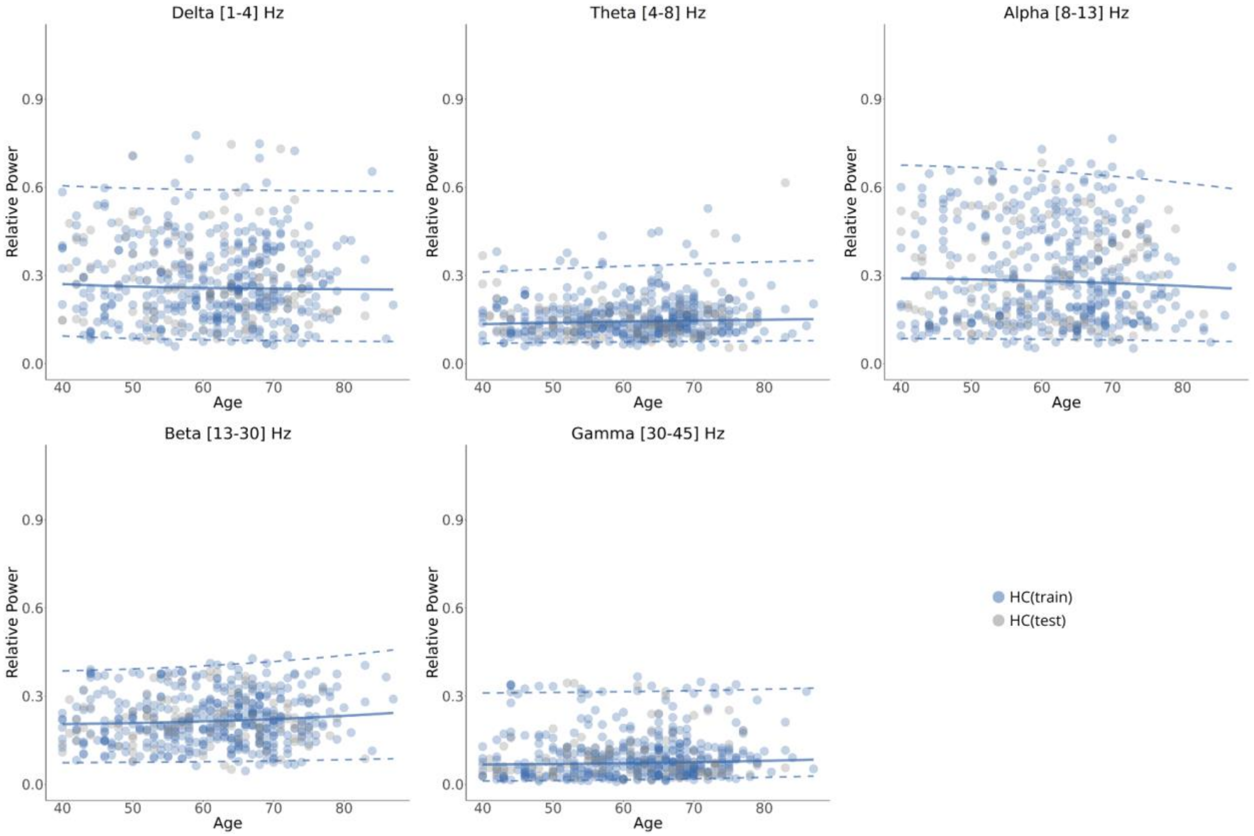


**Fig. S7 | Normative model aging trajectories of spectral features across age within each frequency band along with patients data.** The median (50th percentile) is depicted with a solid blue line, while the 5th and 95th percentiles are indicated by dotted blue lines. The solid black line represents the mean parameter of the GAMLSS fit model. Relative power averaged over channels are represented by scatter points for healthy and patient groups (HC(train), HC(test).

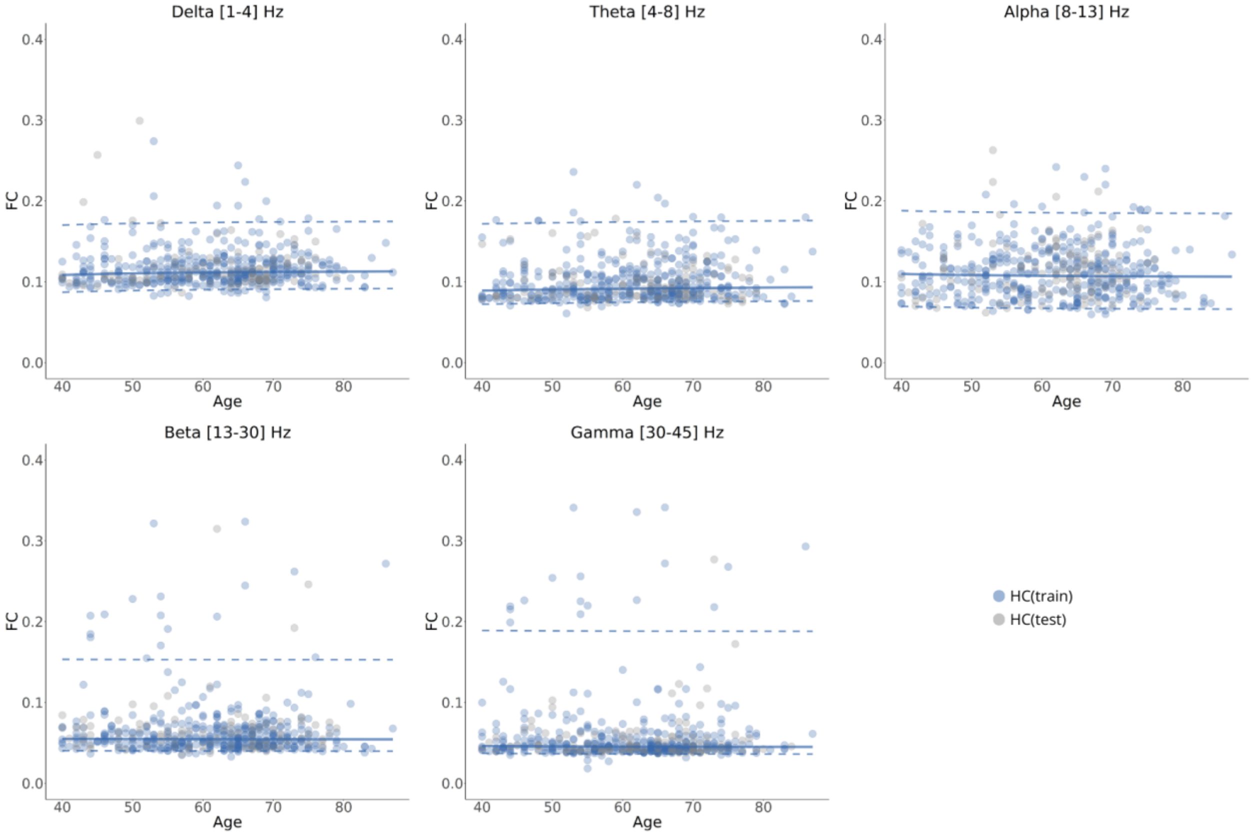


**Fig. S8 | Normative model aging trajectories of functional connectivity features across age within each frequency band along with patients data.** The median (50th percentile) is depicted with a solid blue line, while the 5th and 95th percentiles are indicated by dotted blue lines. The solid black line represents the mean parameter of the GAMLSS fit model. FC values averaged over connections are represented by scatter points for healthy and patient groups (HC(train), HC(test).

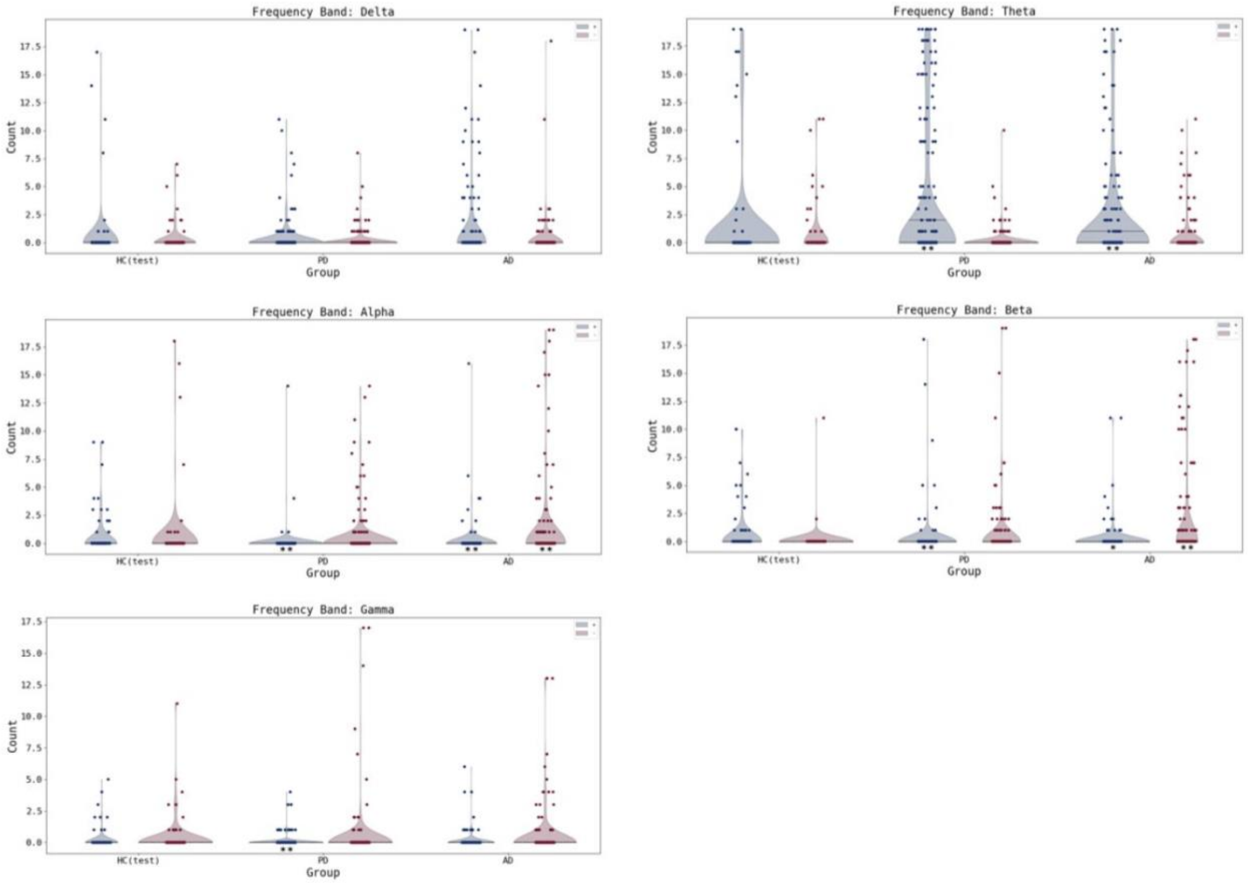


**Fig. S9 | Distribution of the number of extremely deviated channels per subject across groups within each frequency band.** Blue (Red) violins represent positive (negative) deviations. (*) denotes significant difference between HC and cases *p<0.05*. (**) denotes significant difference between HC and cases *p<0.01*.

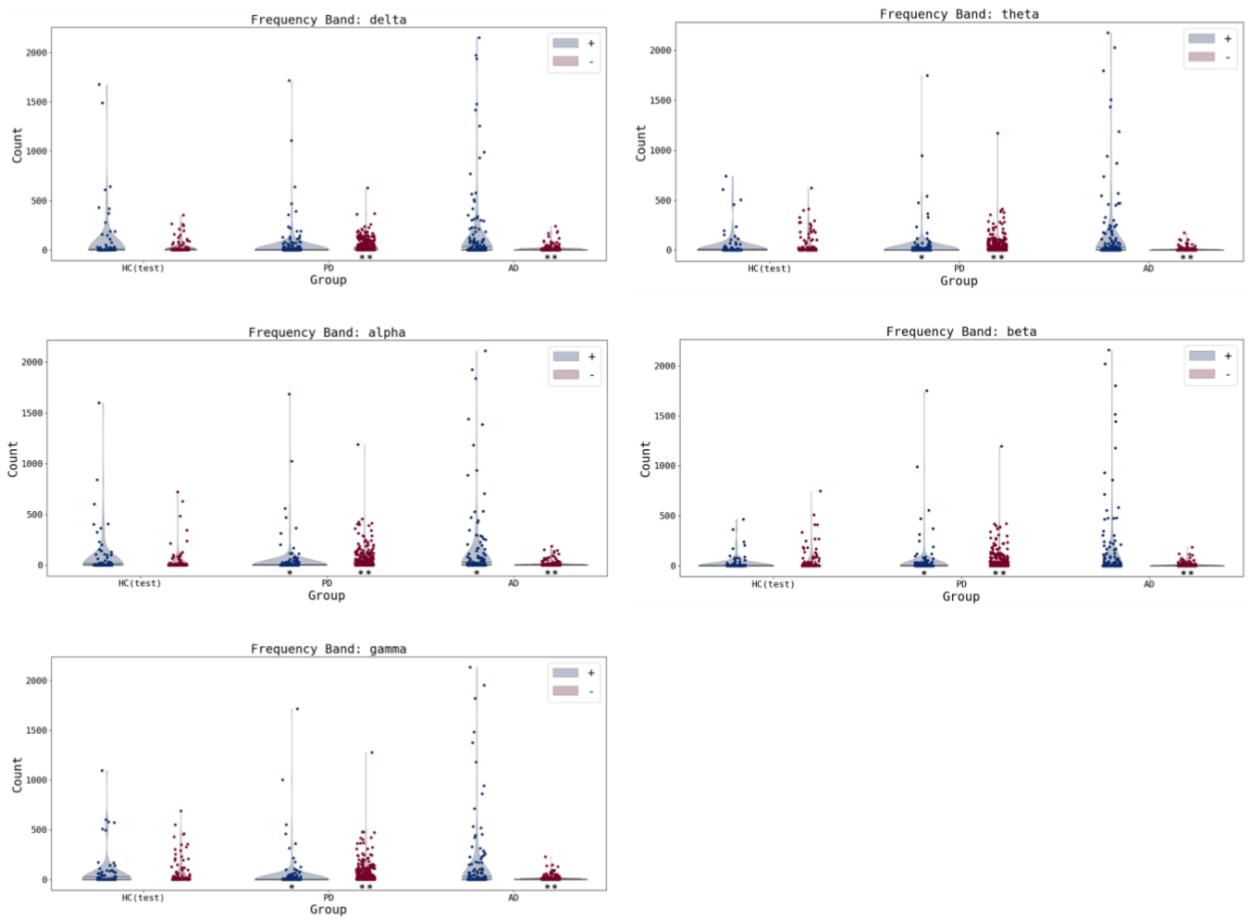


**Fig. S10 | Distribution of the number of extremely deviated connections per subject across groups within each frequency band.** Blue (Red) violins represent positive (negative) deviations. (*) denotes significant difference between HC and cases *p<0.05*. (**) denotes significant difference between HC and cases *p<0.01*.

**Table. S6 | Percentage of subjects exhibiting at least one extremely deviant channel and the median number of extreme deviations across groups within each frequency band.**

| Group | % at least one positive deviation | Median [range] positive deviation | % at least one negative deviation | Median [range] negative deviation |
|---|---|---|---|---|
| **Delta** | | | | |
| HC(test) | 8.08 | 0.0 [0-17] | 10.10 | 0.0 [0-7] |
| PD | 13.14 | 0.0 [0-11] | 8.47 | 0.0 [0-8] |
| AD | 17.77 | 0.0 [0-19] | 9.14 | 0.0 [0-18] |
| **Theta** | | | | |
| HC(test) | 13.27 | 0.0 [0-19] | 14.29 | 0.0 [0-11] |
| PD | 31.36 | 0.0 [0-19] | 7.63 | 0.0 [0-10] |
| AD | 27.41 | 0.0 [0-19] | 13.71 | 0.0 [0-11] |
| **Alpha** | | | | |
| HC(test) | 15.15 | 0.0 [0-9] | 9.09 | 0.0 [0-18] |
| PD | 1.69 | 0.0 [0-14] | 11.86 | 0.0 [0-14] |
| AD | 4.57 | 0.0 [0-16] | 18.78 | 0.0 [0-19] |
| **Beta** | | | | |
| HC(test) | 16.16 | 0.0 [0-10] | 2.02 | 0.0 [0-11] |
| PD | 4.66 | 0.0 [0-18] | 12.71 | 0.0 [0-19] |
| AD | 6.60 | 0.0 [0-11] | 23.35 | 0.0 [0-18] |
| **Gamma** | | | | |
| HC(test) | 9.00 | 0.0 [0-18] | 12.00 | 0.0 [0-10] |
| PD | 5.51 | 0.0 [0-6] | 6.36 | 0.0 [0-18] |
| AD | 7.61 | 0.0 [0-9] | 12.18 | 0.0 [0-16] |

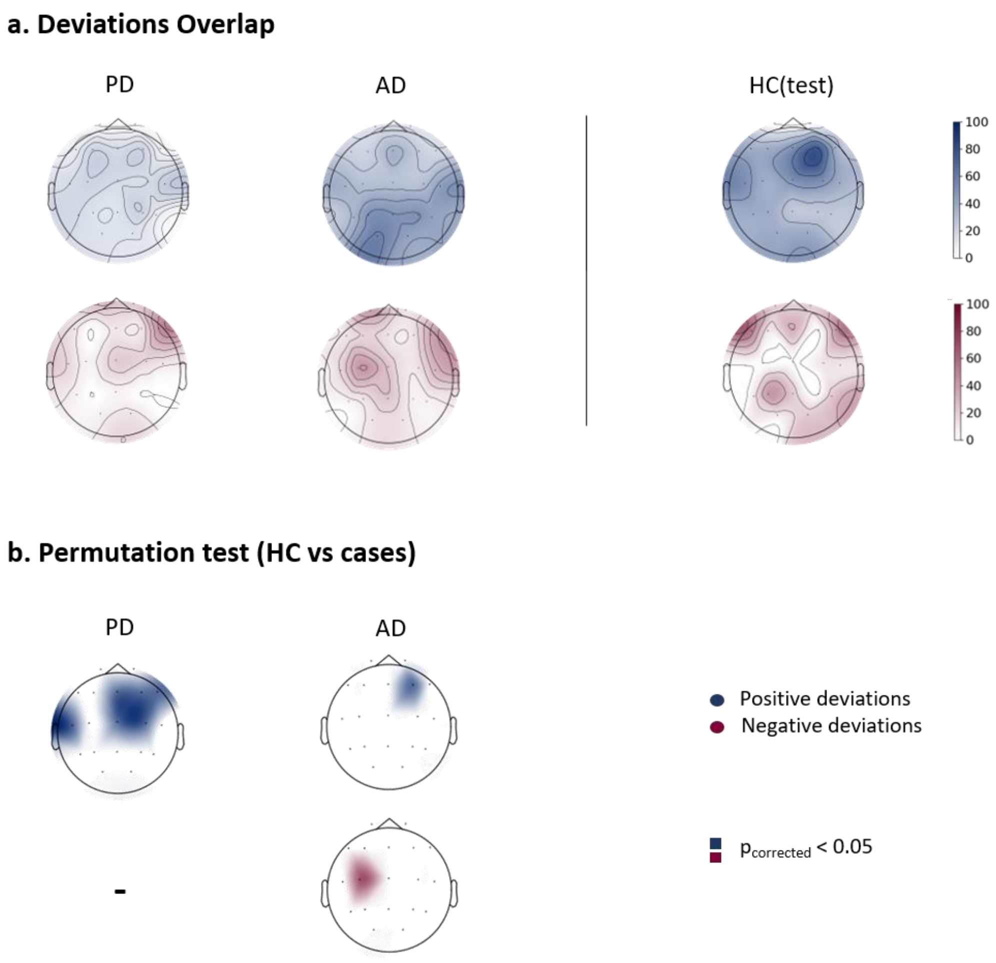


**Fig. S11 | Spectral features maps in the delta band.** (a) Overlap maps of deviation scores for clinical groups and the held-out healthy control group (HC(test)), illustrating areas of common deviation. (b) channels showing significant differences between HC(test) and clinical groups, determined through group-based permutation tests ($p<0.05$, FDR corrected).

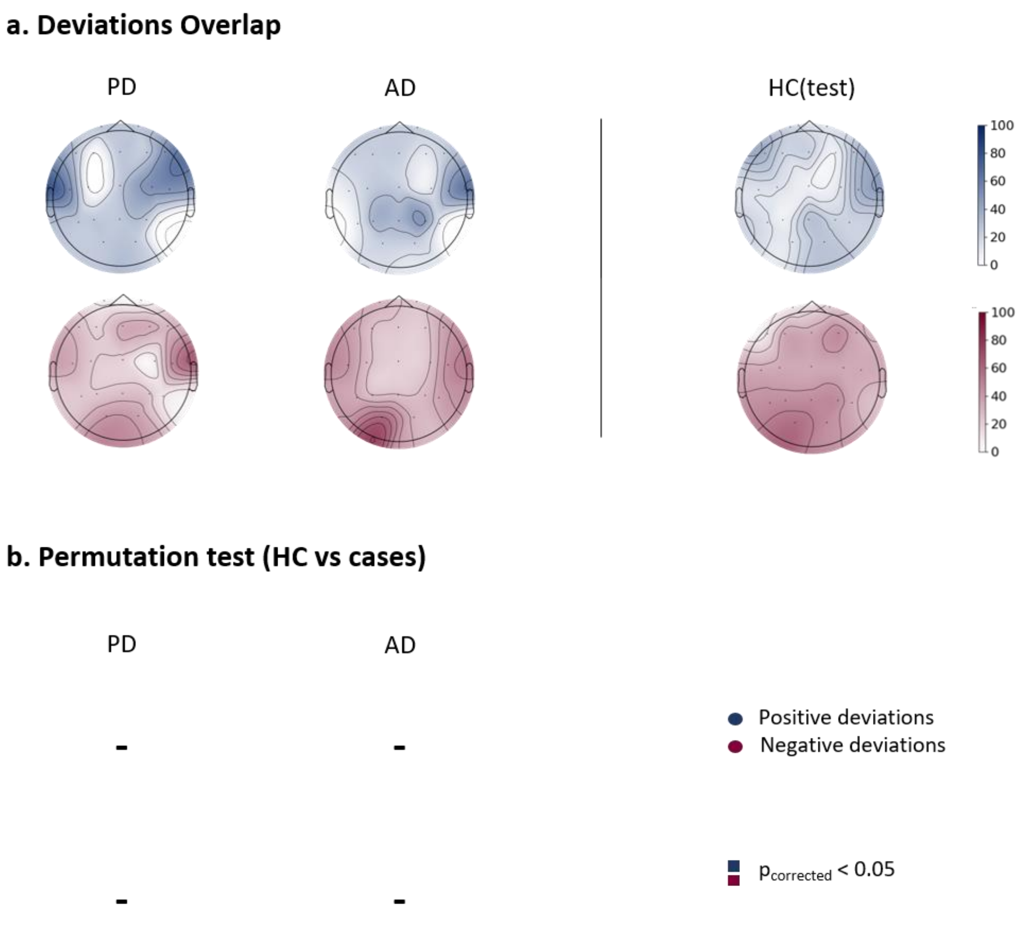


**Fig. S12 | Spectral features maps in the alpha band.** (a) Overlap maps of deviation scores for clinical groups and the held-out healthy control group (HC(test)), illustrating areas of common deviation. (b) channels showing significant differences between HC(test) and clinical groups, determined through group-based permutation tests ($p<0.05$, FDR corrected).

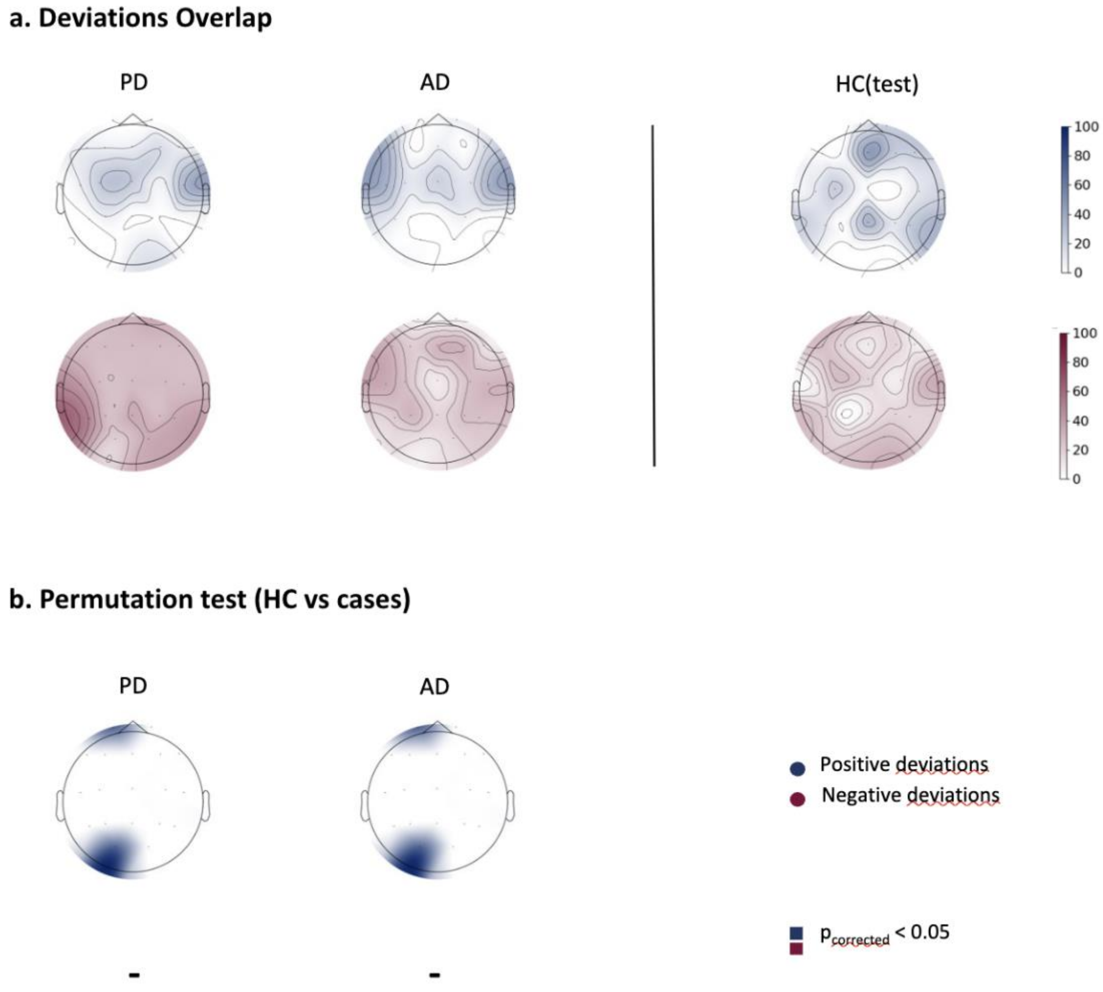


**Fig. S13 | Spectral features maps in the gamma band.** (a) Overlap maps of deviation scores for clinical groups and the held-out healthy control group (HC(test)), illustrating areas of common deviation. (b) channels showing significant differences between HC(test) and clinical groups, determined through group-based permutation tests (p<0.05, FDR corrected).

**Table. S7 | Percentage of subjects exhibiting at least one extremely deviant connection and the median number of extreme deviations across groups within each frequency band.**

| Group | % at least one positive deviation | Median [range] positive deviation | % at least one negative deviation | Median [range] negative deviation |
|---|---|---|---|---|
| **Delta** | | | | |
| HC(test) | 39.80 | 0.0 [0-1676] | 72.45 | 3.0 [0-353] |
| PD | 40.68 | 0.0 [0-1716] | 86.86 | 26.0 [0-627] |
| AD | 45.69 | 0.0 [0-2150] | 53.81 | 1.0 [0-240] |
| **Theta** | | | | |
| HC(test) | 42.42 | 0.0 [0-740] | 70.71 | 4.0 [0-621] |
| PD | 35.17 | 0.0 [0-1747] | 83.47 | 17.5 [0-1170] |
| AD | 46.70 | 0.0 [0-2175] | 49.24 | 0.0 [0-173] |
| **Alpha** | | | | |
| HC(test) | 45.92 | 0.0 [0-1598] | 54.08 | 1.0 [0-721] |
| PD | 38.14 | 0.0 [0-1684] | 83.05 | 19.0 [0-1187] |
| AD | 52.79 | 1.0 [0-2110] | 53.30 | 1.0 [0-185] |
| **Beta** | | | | |
| HC(test) | 42.27 | 0.0 [0-464] | 62.89 | 2.0 [0-747] |
| PD | 37.71 | 0.0 [0-1750] | 83.05 | 17.0 [0-1194] |
| AD | 46.70 | 0.0 [0-2157] | 49.75 | 0.0 [0-185] |
| **Gamma** | | | | |
| HC(test) | 42.86 | 0.0 [0-1095] | 73.47 | 7.0 [0-690] |
| PD | 37.29 | 0.0 [0-1716] | 85.59 | 26.0 [0-1277] |
| AD | 44.16 | 0.0 [0-2134] | 54.82 | 1.0 [0-228] |

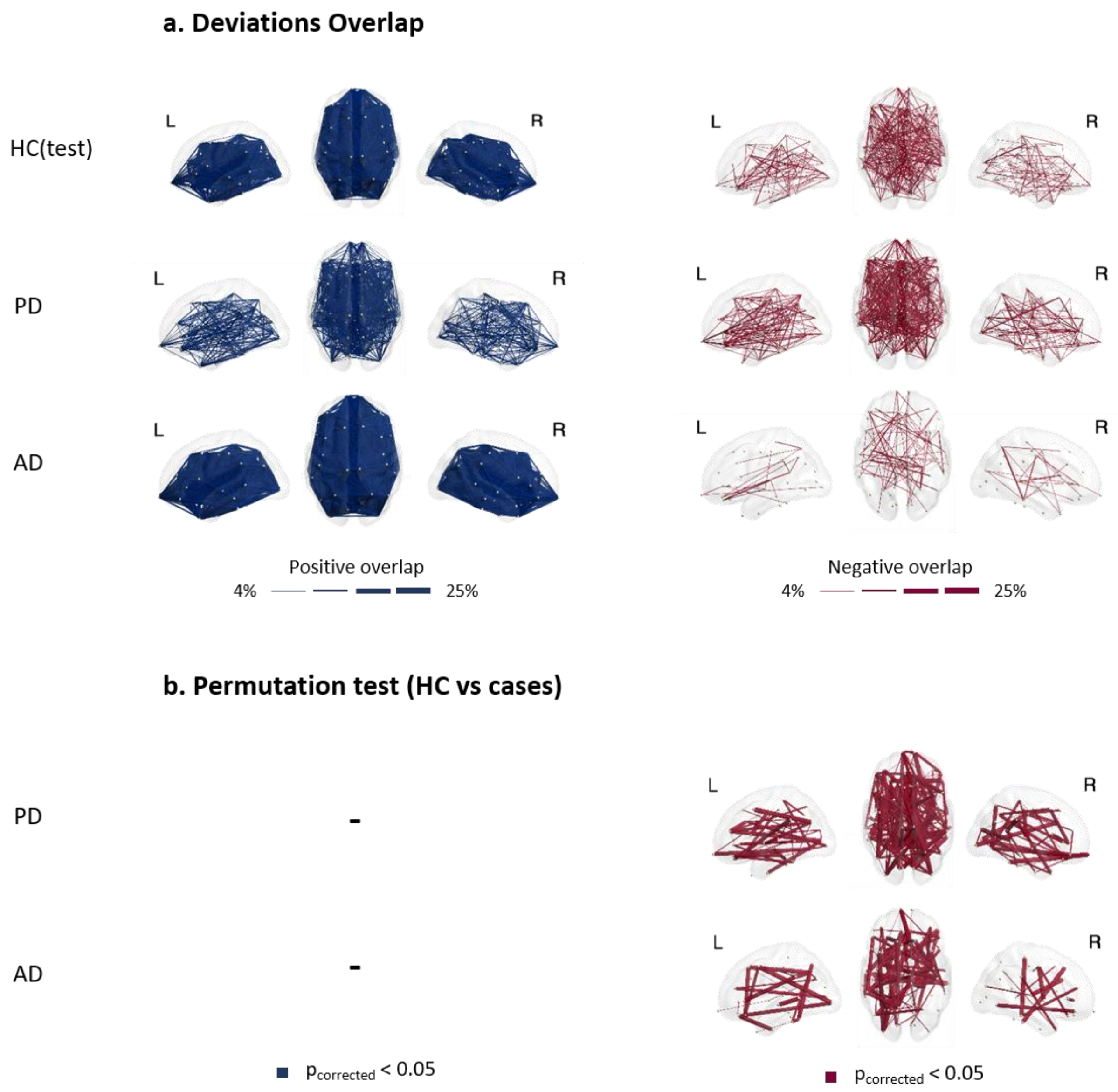


**Fig. S14 | Functional connectivity maps in delta band. (a)** Overlap maps of positive and negative deviation scores for clinical groups and the held-out healthy control group (HC(test)), illustrating areas of common deviation among patients (with only the highest 4% overlap values being plotted for visualization purposes). **(b)** functional connections showing significant differences between HC(test) and clinical groups, determined through group-based permutation tests (p<0.05, FDR corrected).

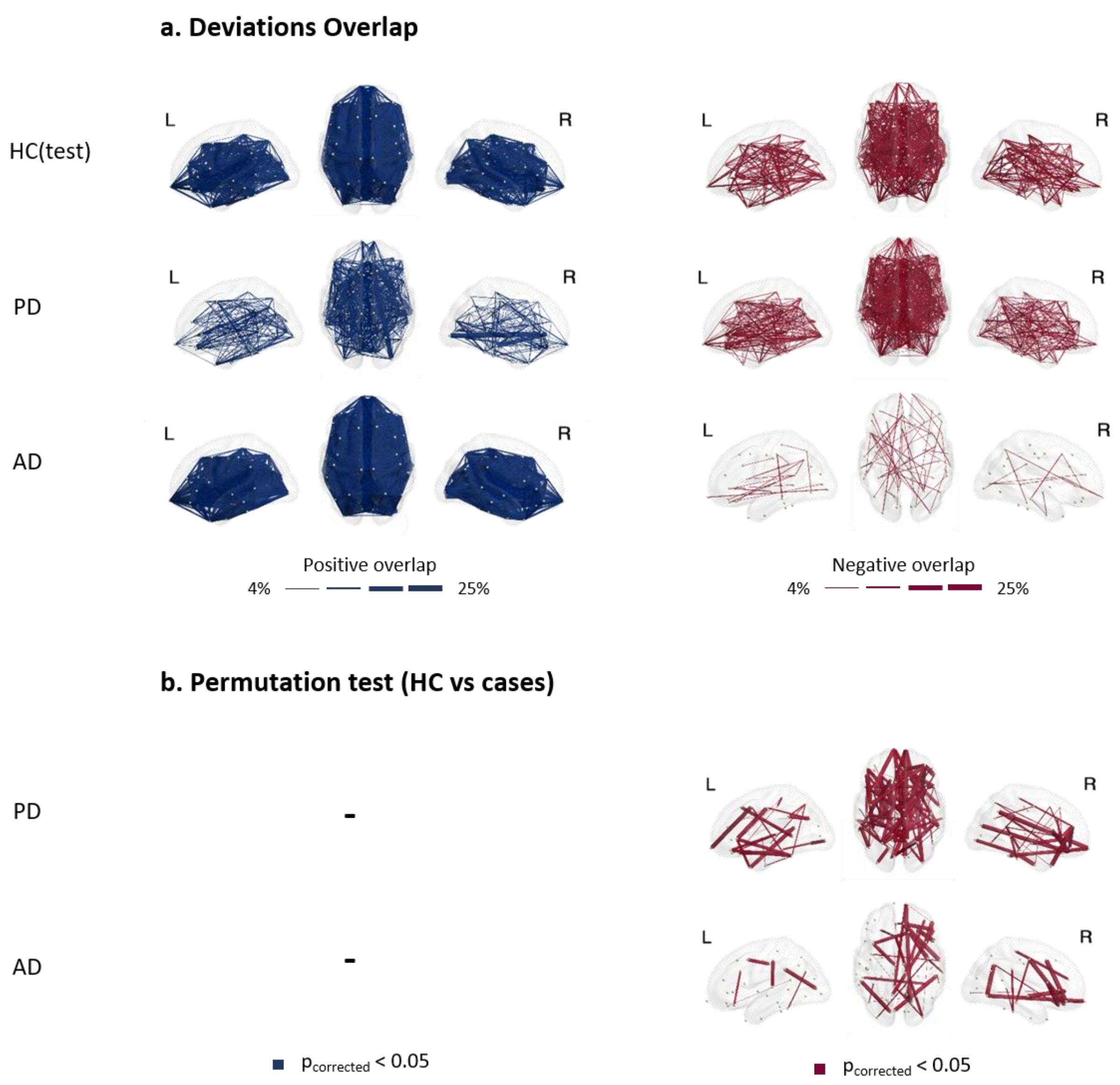


**Fig. S15 | Functional connectivity maps in alpha band. (a)** Overlap maps of positive and negative deviation scores for clinical groups and the held-out healthy control group (HC(test)), illustrating areas of common deviation among patients (with only the highest 4% overlap values being plotted for visualization purposes). **(b)** functional connections showing significant differences between HC(test) and clinical groups, determined through group-based permutation tests ($p<0.05$, FDR corrected).

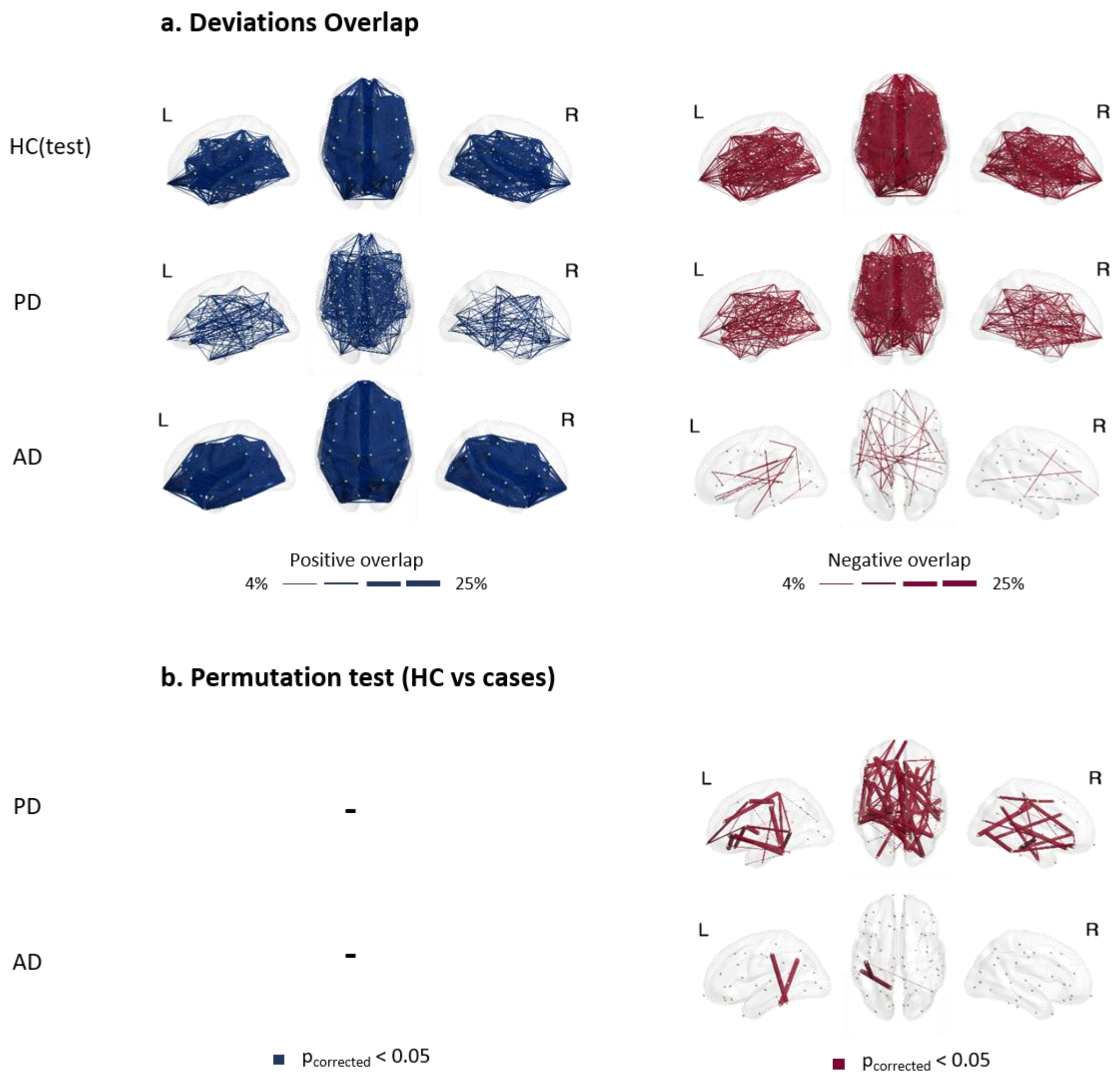


**Fig. S16 | Functional connectivity maps in gamma band. (a)** Overlap maps of positive and negative deviation scores for clinical groups and the held-out healthy control group (HC(test)), illustrating areas of common deviation among patients (with only the highest 4% overlap values being plotted for visualization purposes). **(b)** functional connections showing significant differences between HC(test) and clinical groups, determined through group-based permutation tests (p<0.05, FDR corrected).

**Table. S8 | Distribution of the proportion of the significant connections (after permutation test) in overlap maps across resting state networks for clinical groups in all frequency bands.** Values in red indicate proportions above 35%. *SAN denotes salience, DMN denotes default mode, VIS denotes visual, TEMP denotes temporal, CCN denotes cognitive control, FPN denotes frontoparietal, AUD denotes auditory, SMN denotes somatomotor, and DAN denotes dorsal attention networks.*

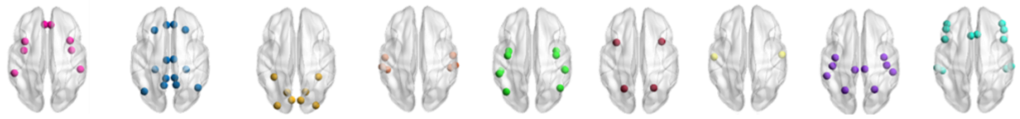

| Prop (%) | | SAN | DMN | VIS | TEMP | CCN | FPN | AUD | SMN | DAN |
|---|---|---|---|---|---|---|---|---|---|---|
| ***Delta*** | **PD** | 23.67 | 39.05 | 24.26 | 13.61 | 29.59 | 10.06 | 27.81 | 3.55 | 33.14 |
| | **AD** | 30.99 | 43.66 | 19.72 | 15.49 | 38.03 | 11.27 | 29.58 | 5.63 | 33.80 |
| ***Theta*** | **PD** | 16.09 | 40.23 | 26.44 | 12.64 | 21.84 | 8.05 | 26.44 | 1.15 | 34.48 |
| | **AD** | 16.67 | 35 | 30 | 18.33 | 31.67 | 10 | 30 | 6.67 | 33.33 |
| ***Alpha*** | **PD** | 17.92 | 41.51 | 20.75 | 14.15 | 21.70 | 15.09 | 24.53 | 3.77 | 35.85 |
| | **AD** | 33.33 | 44.44 | 33.33 | 19.44 | 36.11 | 11.11 | 22.22 | 5.55 | 36.11 |
| ***Beta*** | **PD** | 14 | 28 | 30 | 16 | 26 | 10 | 40 | 10 | 30 |
| | **AD** | 21.13 | 26.76 | 22.54 | 21.13 | 19.72 | 9.86 | 28.17 | 7.04 | 40.85 |
| ***Gamma*** | **PD** | 28.40 | 33.33 | 16.05 | 19.75 | 24.69 | 13.58 | 34.57 | 7.41 | 37.04 |
| | **AD** | 0 | 66.67 | 33.33 | 33.33 | 66.67 | 0 | 66.67 | 0 | 33.33 |

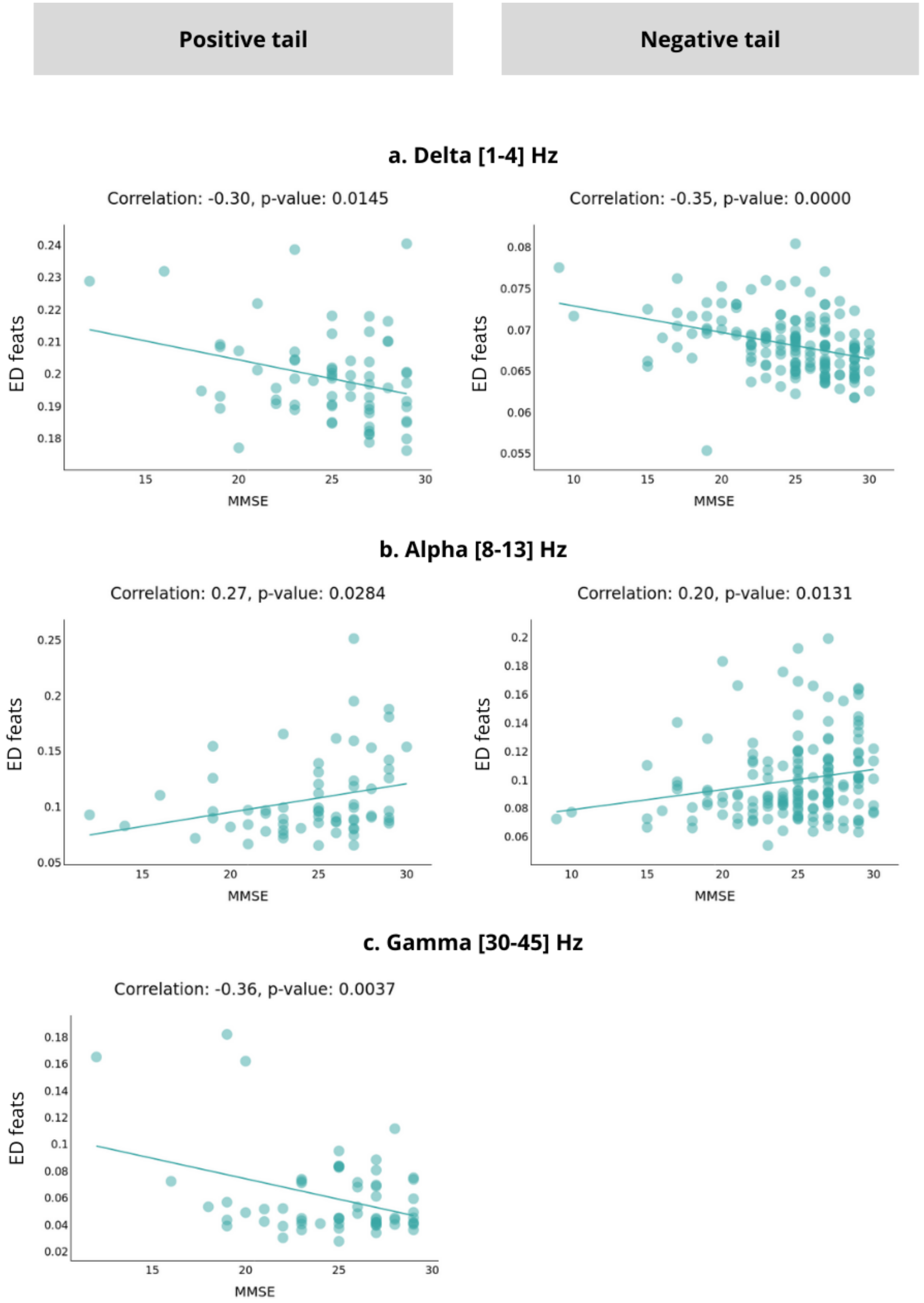

Positive tail
Negative tail
a. Delta [1-4] Hz
Correlation: -0.30, p-value: 0.0145
Correlation: -0.35, p-value: 0.0000
ED feats
MMSE
b. Alpha [8-13] Hz
Correlation: 0.27, p-value: 0.0284
Correlation: 0.20, p-value: 0.0131
ED feats
MMSE
c. Gamma [30-45] Hz
Correlation: -0.36, p-value: 0.0037
ED feats
MMSE

**Fig. S17 | Correlations between PD subjects' scores and clinical assessment scores (MMSE) (p-value<0.05) in delta, alpha, and gamma bands respectively. (a, b, and c)** Scores are computed as the averaged FC features over the extremely deviated connections (positive and negative). The corresponding correlation coefficient and p-value are indicated for each case.

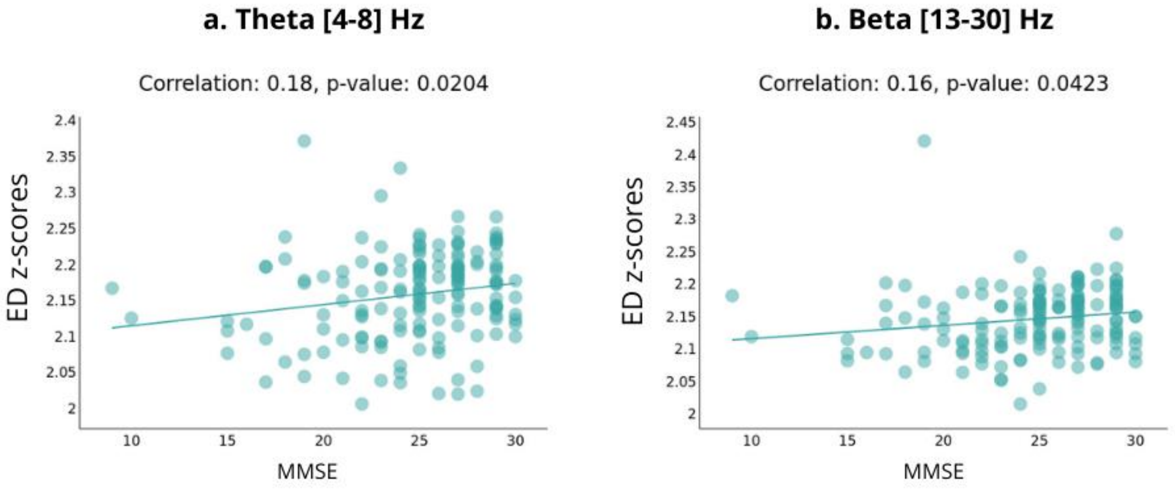


**Fig. S18 | Significant correlation between PD subjects' scores and clinical assessment scores (MMSE) (p-value<0.05) in theta, and beta bands respectively. (a, b)** Scores are computed as the averaged z-scores over the extremely deviated connections for FC features. The corresponding correlation coefficient and p-value are indicated for each case.
